# Virtue as the mean: Pan-human consensus genome significantly improves the accuracy of RNA-seq analyses

**DOI:** 10.1101/2020.12.22.423111

**Authors:** Benjamin Kaminow, Sara Ballouz, Jesse Gillis, Alexander Dobin

## Abstract

The Human Reference Genome serves as the foundation for modern genomic analyses. However, in its present form, it does not adequately represent the vast genetic diversity of the human population. In this study, we explored the consensus genome as a potential successor of the current Reference genome and assessed its effect on the accuracy of RNA-seq read alignment. In order to find the best haploid genome representation, we constructed consensus genomes at the Pan-human, Super-population and Population levels, utilizing variant information from the 1000 Genomes project. Using personal haploid genomes as the ground truth, we compared mapping errors for real RNA-seq reads aligned to the consensus genomes versus the Reference genome. For reads overlapping homozygous variants, we found that the mapping error decreased by a factor of ~2-3 when the Reference was replaced with the Pan-human consensus genome. Interestingly, we also found that using more population-specific consensuses resulted in little to no increase over using the Pan-human consensus, suggesting a limit in the utility of incorporating more specific genomic variation. To assess the functional impact, we compared splice junction expression in the different genomes and found that the Pan-human consensus increases accuracy of splice junction quantification for hundreds of splice junctions.

## Background

In 2003, 15 years of work culminated with the International Human Genome Sequencing Consortium publishing the first finished version of the Human Reference Genome (https://www.genome.gov/human-genome-project/Completion-FAQ; IHGSC 2004). Despite the utility and continuous improvements over the years, it is still not without flaws – primarily the lack of variation information. Around 93% of the current GRCh38 assembly is composed of DNA from just 11 individuals (https://www.ncbi.nlm.nih.gov/grc/help/faq/; Lander et al. 2001). Because such a large portion of the Reference comes from such a small pool of individuals, it does not adequately represent the vast diversity present in the human population (Chen and Butte 2011; Rosenfeld et al. 2012; Sherman et al. 2019). To explore and capture human diversity, researchers have continued sequencing thousands of genomes. The first of such projects, the 1000 Genomes Project, sequenced 2,504 individuals across 26 populations. Overall, it was estimated that ~3,000 genomes would be necessary to capture the most common variants (Ionita-Laza et al. 2009), while structural variation present in the human populations has challenged this (Berlin et al. 2015). One particularly glaring example was shown in a recent construction of an African pan-genome, which contained almost 300M bases of DNA not seen in GRCh38 (Sherman et al. 2019). This lack of variation information negatively affects all kinds of genomic analyses that utilize the Reference, such as disease studies and GWAS analyses (Buchkovich et al. 2015; Castel et al. 2015; Chen and Butte 2011; Rosenfeld et al. 2012; Sherman et al. 2019; Stevenson et al. 2013). However, despite the ubiquity of RNA-seq alignment and quantification, the improvements on mapping from using a more diverse reference have not been shown.

While graph genomes are theoretically capable of encapsulating all observed variation information (Church et al. 2015; Garrison et al. 2018; Paten et al. 2017; Rakocevic et al. 2019; Sirèn et al. 2020; Valenzuela et al. 2018), it remains difficult to use these tools for large scale expression analysis such as in RNA-seq quantification. In prior work, we proposed the use of a consensus genome to inherently capture common variation, whilst still retaining the structure and functionality of the current Reference assembly (Ballouz et al. 2019). A consensus genome is a linear haploid genome that incorporates population variation information by replacing all minor alleles in the Reference genome with the major allele of that variant (Balasubramanian et al. 2011; Ballouz et al. 2019; Barbitoff et al. 2018; Dewey et al. 2011; Karthikeyan et al. 2016; Pritt et al. 2018; Shukla et al. 2019) (Figure 1a). Because allele frequencies must be defined with respect to a population, a consensus genome is representative of the population used to define the major and minor alleles. Prior work has shown that using a consensus genome can have positive effects on variant calling, (Karthikeyan et al. 2016; Pritt et al. 2018; Shukla et al. 2019) and construction of population-specific consensus genomes has been a major goal of multiple projects (Cho et al. 2016; Fakhro et al. 2016; Higasa et al. 2016; Sherman et al. 2019; Takayama et al. 2019). Additionally, replacing the current Reference genome with a consensus genome in existing analysis pipelines is straightforward, since the consensus genome is still a linear haploid sequence.

**Figure 1:**
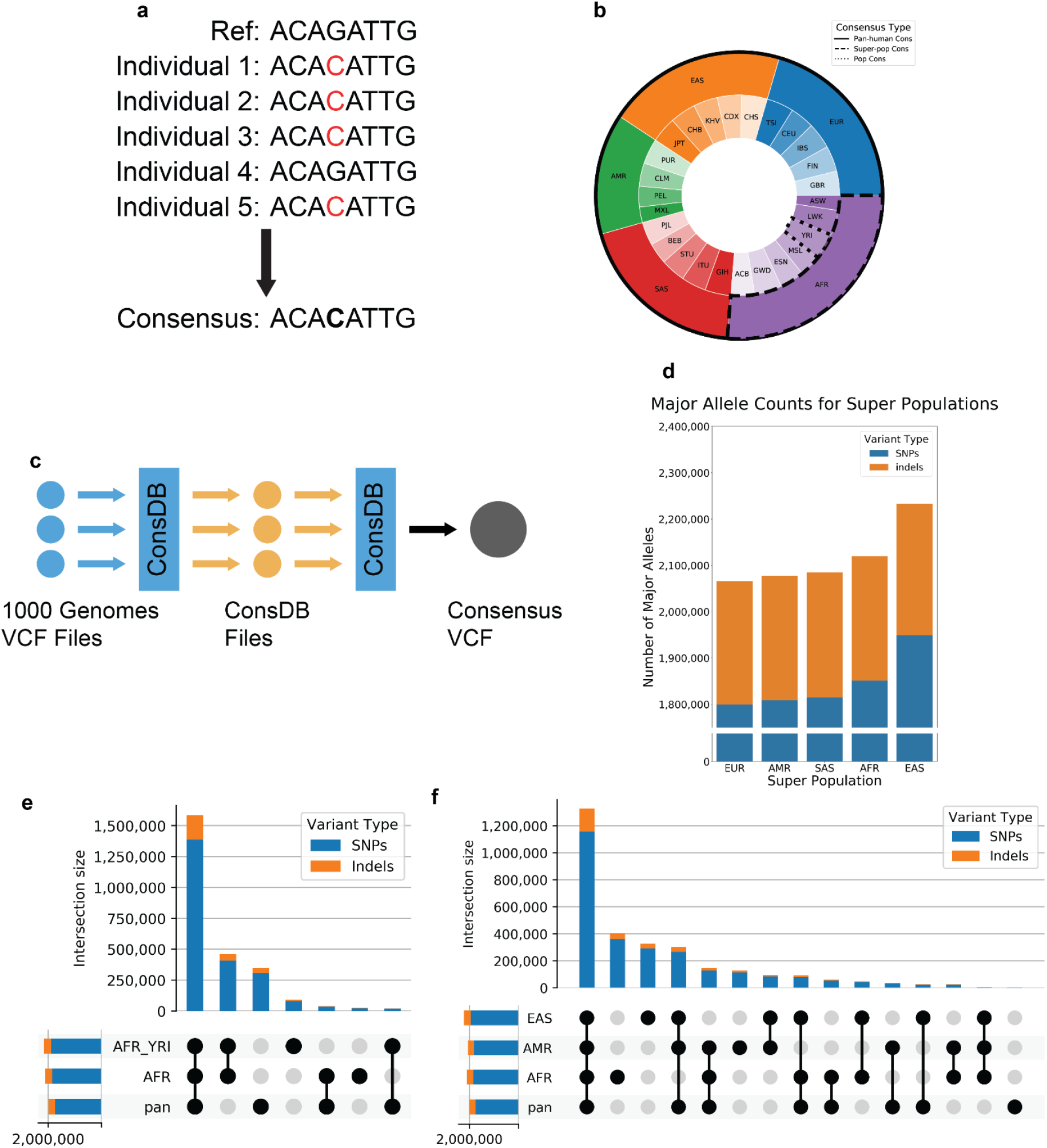
**a)** Construction of a consensus genome; the minor allele in the Reference is replaced by the most common allele in the population. **b)** Visual representation of the individuals used to construct consensus genomes of varying population specificity. **c)** ConsDB workflow. **d)** Number of major alleles for each population consensus genome that were replaced in the Reference. **e)** Number of SNPs and indels shared between different combinations of the Pan-human, Super-population, and Population consensus genomes for the African population. The bars in the top bar plot show the number of SNPs and indels that are unique to the intersection of genomes indicated in the dot matrix below. The horizontal bars on the bottom left show the total number of SNPs and indels present in each genome. **f)** Number of SNPs and indels shared between different combinations of the Pan-human consensus and all 3 super-population consensus genomes. The bars in the top bar plot show the number of SNPs and indels that are unique to the intersection of genomes indicated in the dot matrix below. The horizontal bars on the bottom left show the total number of SNPs and indels present in each genome.

Here, we seek to answer the question of which linear reference representation is best for RNA-seq mapping and downstream analyses. We considered several consensus genomes, built by replacing all minor alleles in the reference with the major alleles at different population levels: pan-human, super-population, and population. To work with consensus genomes, we developed ConsDB to construct pan-human and population-level consensuses, and STAR-consensus to streamline RNA-seq mapping to consensus genomes. We defined the ground truth by mapping the individuals’ RNA-seq reads to their own personal haploid genomes, and evaluated the mapping accuracy improvements arising from replacing the GRCh38 reference with the Pan-human consensus, Super-population or Population consensus genomes. We found that for all individuals, the Pan-human consensus decreased the mapping error from the Reference by ~2-3 fold, while the Super-population and Population consensuses did not perform significantly better than the Pan-human consensus. To assess the functional impact, we measured errors in splice junction expression quantification for different genome representations with respect to the ground truth of the personal genome. We again found that the Pan-human consensus offers an improvement over the Reference, with ~5 times as many splice junctions having a larger quantification error for the Reference than for the Pan-human consensus.

## Pan-human Consensus captures the majority of population deviation from the Reference

The construction of consensus genomes requires population allele frequency information. Currently, several databases exist that contain this information (Auton et al. 2015; Church et al. 2015; Karczewski et al. 2020; Sherry et al. 2001). In this study we utilized the 1000 Genomes Project database, which was established in order to discover and catalogue human genome variant information (Auton et al. 2015; Clarke et al. 2017). In order to avoid population bias, the individuals genotyped in the 1000 Genomes Project were selected to create an even population distribution across 26 populations, which are grouped into 5 super-populations (Auton et al. 2015) (Figure 1b). The information from the 1000 Genomes Project is available through the International Genome Sample Resource (IGSR), and can be downloaded in the form of VCF files, which contain variant genotype information for all of the individuals contained in the analysis (Auton et al. 2015).

We constructed three types of consensus genomes based on the various population levels present in the 1000 Genomes Project: a Pan-human consensus genome, a Super-population consensus genome, and a Population consensus genome (Figure 1b). For the Pan-human consensus we calculated allele frequency using genotype information from all individuals present in the database. For the Super-population and Population consensuses, we used genotype information from all individuals of a given super-population or population. For the 8 individuals whose RNA-seq data we utilized in this study, we used the consensus genomes built from the super-population and population to which each individual belongs. To construct these consensuses, we replaced all minor alleles (alleles with a population allele frequency AF < 0.5) present in the Reference with the major alleles (AF > 0.5). We will call these variants replaced in the reference the major allele replacements (MAR). The release of the 1000 Genomes database that we used contained only biallelic variants, i.e. each variant had exactly one minor allele and one major allele. Additionally, it only contained SNPs and small insertions and deletions (<50 bp), while large structural variants were not considered in this study. Although SVs are a large source of genomic variation, they are understudied and not sufficiently catalogued to be used in consensus genomes due to mapping and classification difficulties (Mahmoud et al. 2019).

In order to facilitate working with the large VCF files of the 1000 Genomes Project database, we developed ConsDB, a Python package that provides a convenient, class-based interface to work with the large number of variants contained in the 1000 Genomes Project database. It also provides a main script with a number of run modes to perform common tasks associated with consensus genomes, such as the construction of the consensus genome VCF files used in this study. ConsDB operates using a simple workflow (Figure 1c). The first step is downloading the database VCF files. For this study, we used the 1000 Genomes Project, but ConsDB is also capable of parsing gnomAD VCF files. The next step is for ConsDB to parse the database VCF files and save them in the ConsDB format. At this point, files from different databases (if multiple databases are being used) can be combined into one file per chromosome. Finally, ConsDB uses these parsed files to generate the end result, in this case a VCF file defining a consensus genome.

The personal haploid genomes were constructed using the individual genotypes from the 1000 Genomes Project database. For each individual, all homozygous variants that differ from the Reference were inserted into the Reference. Additionally, all heterozygous alleles were randomly chosen with a probability of 0.5 to be included or excluded. Although these haploid personal genomes are a crude approximation of the true diploid genome, they are sufficient for comparison of mapping accuracy between haploid consensuses and the haploid Reference, and thus we used them to define the ground truth for RNA-seq mapping in this study.

Figure 1d shows the number of minor alleles in the GRCh38 reference that have to be replaced with the major alleles for each of the Super-population consensus genomes. The European consensus is the most similar to the Reference, and it still requires ~2.1 million SNP and indel corrections from the Reference. Other Super-population consensuses contain even larger numbers of major allele deviations from the Reference, with the East Asian consensus differing most from the Reference. We note that such a large number of minor alleles in the Reference with respect to any population stems from its construction, which utilized sequences from only one individual for most of the genomic loci, and thus incorporated individual-specific low frequency alleles.

In Figure 1e, we compute intersections of the MARs in the Pan-human, African Super-population, and Yoruban Population consensus genomes. The Pan-human consensus shares most of the major alleles with the Super- and Population consensuses (~1.5M), while the latter two share ~400k MARs not present in the Pan-human consensus. The Pan-human consensus contains ~300k MARs not present in either Super- or Population consensuses. Finally, the Yoruban Population consensus contains ~50k unique MARs. The intersections of MARs look similar for other populations (Supplementary Figures S1-2) as well as personal homozygous variants (Supplementary Figures S3-5). Figure 1f shows the intersections between the MARs for the Pan-human consensus and 3 Super-population consensuses. The MARs shared by all four of these genomes make up the largest group, which contains ~1.2M MARs and represents well over half of the MARs in any one genome. This group is more than 3 times as large as the next largest group, again demonstrating that the majority of the population deviation from the Reference is captured in the Pan-human consensus.

## Consensus genomes significantly improve RNA-seq mapping

Next, we analyzed to what extent the consensus genomes improve RNA-seq mapping accuracy. The RNA-seq reads were taken from the Human Genome Structural Variation Consortium, which sequenced three father-mother-daughter trios from the 1000 Genomes Project (Fairley et al. 2020). One of these individuals (HG00514 from the East Asian trio) is not present in the database version used in this analysis and was excluded from our analysis.

To simplify alignment to the consensus genome, we developed STAR-consensus, an extension to the RNA-seq aligner STAR (Figure 2a) (Dobin et al. 2013). It imports variants from a VCF file and incorporates them into the reference genome sequence, thus creating a transformed genome for mapping. Importantly, after mapping the reads to the transformed genome, STAR-consensus can perform a reverse transformation of the alignment coordinates back to the original reference genome coordinates. This transformation is non-trivial when insertion or deletion variants are included, and allows performing all downstream analyses in the reference coordinate system. Such an approach is an incremental but important step towards taking advantage of the consensus genome, while at the same time utilizing the conventional coordinate system.

**Figure 2:**
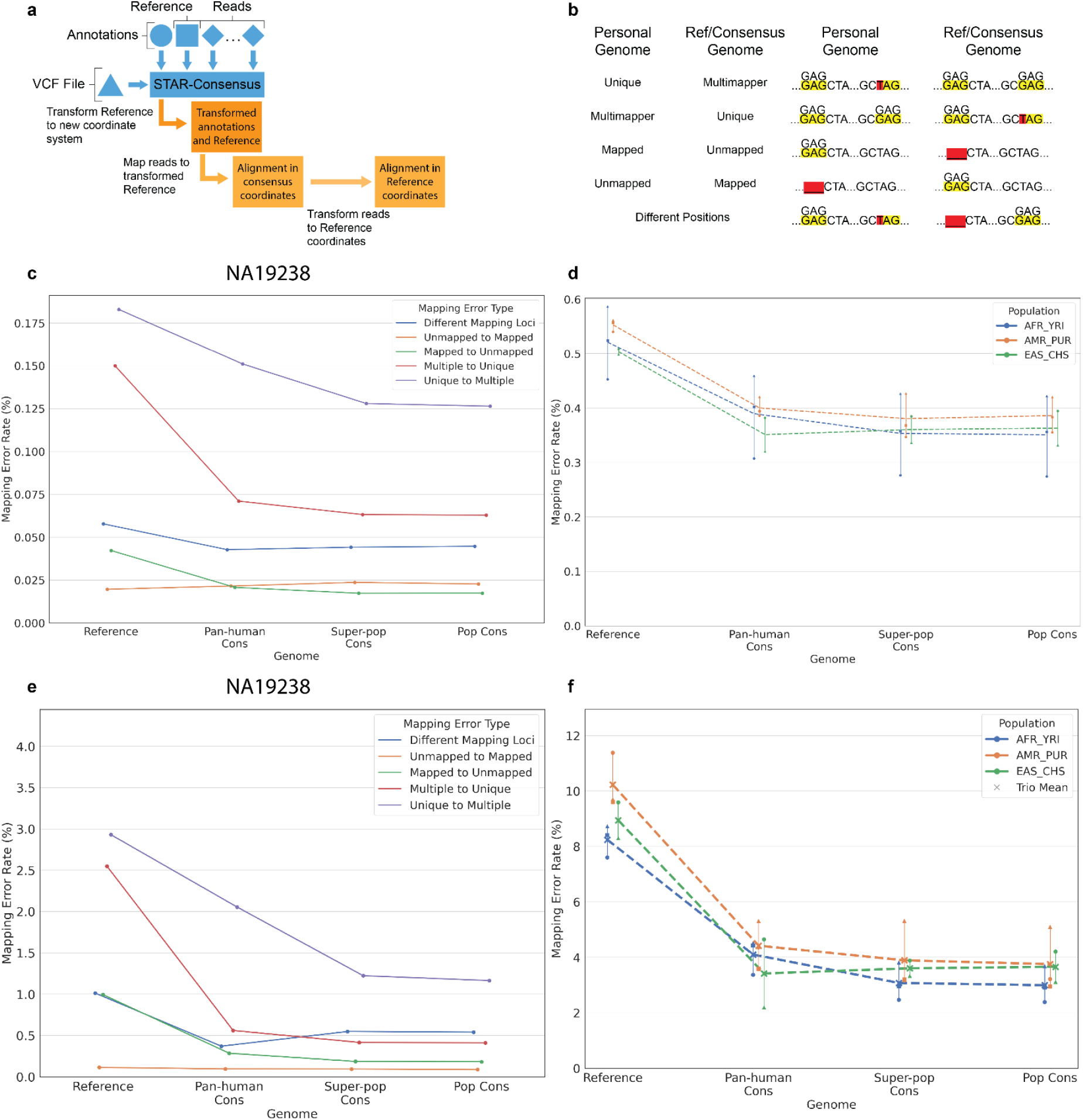
**a)** Internal workflow of STAR-consensus. **b)** Different types of mapping errors based on the read’s mapping status in the individual’s haploid personal genome and the Reference or given consensus genome. **c)** Overall mapping error rate for each error type for individual NA19238. Genome is shown on the x-axis and the mapping error rate is shown on the y-axis. **d)** Overall mapping error rate for all individuals. Genome is shown on the x-axis and the mapping error rate is shown on the y-axis. Individuals from the same population are grouped together by color, with each marker shape representing one individual in the population. The dashed line shows the average error rate for the population and the solid vertical line shows the range of the population. **e)** Homozygous mapping error rate for each error type for individual NA19238. Genome is shown on the x-axis and the mapping error rate is shown on the y-axis. **f)** Homozygous mapping error rate for all individuals. Genome is shown on the x-axis and the mapping error rate is shown on the y-axis. Individuals from the same population are grouped together by color, with each marker shape representing one individual in the population. The dashed line shows the average error rate for the population and the solid vertical line shows the range of the population.

In order to assess error rate, we needed to compare the read mappings in the various genomes to a ground truth. However, because the true mapping location of these reads is unknown, we used the read mappings to the personal haploid genomes as the ground truth. The personal haploid genomes are a close approximation of the true genomes, and therefore the locations to which the reads map in the personal genomes should be quite similar to their true original locations.

We classified mapping errors into five types of errors based on the change of the read’s alignment status in the Reference/consensus genome compared to the ground truth (Figure 2b). The different error types are: reads that are mapped uniquely in the personal genome but mapped to multiple places in the other genome (Unique to Multiple), reads that are mapped to multiple places in the personal genome but mapped uniquely in the other genome (Multiple to Unique), reads that mapped to the personal genome but not to the other genome (Mapped to Unmapped), reads that didn’t map to the personal genome but did map to the other genome (Unmapped to Mapped), and reads that mapped uniquely in both genomes but to different positions (Different Mapping Loci). The mapping error rate for an error type is defined as the number of erroneously mapped reads normalized by the total number of reads from an individual.

For each individual, we calculated the error rates for mapping to the Reference and their respective consensus genomes (Pan-human, Super-population, Population). Figure 2c shows the overall error rates for each error type for the individual NA19238. The largest error comes from the reads that switch from mapping uniquely in the personal genome to mapping to multiple loci in the Reference/consensus genomes, followed by reads that map to multiple loci in the personal genome but map uniquely in the Reference/consensus.

We also separately plotted the error rate for reads that overlap indel variants (Supplementary Figure S6), which are very small compared to the overall error rates in Figure 2c. These plots look similar for the other individuals (Supplementary Figures S7-20).

Figure 2d shows the overall mapping error rate for all eight individuals, summed over the five error types. We see a noticeable decrease in the error rate when the Reference genome is replaced with the Pan-human consensus. Notably, increasing population specificity to the Super-population or Population consensus does not result in a significant further reduction of the error rate. This trend mirrors the observation about the minor alleles discussed above (Figure 1e-f), and supports the conjecture that the majority of the mapping accuracy improvement is captured by the Pan-human consensus, with little additional benefit from the Super-population or Population consensuses.

Replacement of the minor alleles in the Reference with the major alleles in the consensus can only correct the mapping errors caused by the homozygous alternative alleles in an individual. Of course, the actual individual genome is diploid and contains millions of heterozygous variants (i.e. both the major and minor alleles are present), which cannot be truthfully represented in a haploid Reference or consensus genome. To elucidate this issue, we defined the *homozygous mapping error rate* as the number of erroneously mapped reads that overlap homozygous variants normalized by the total number of reads overlapping homozygous variants for an individual. The homozygous mapping error rate shows the effect of different genomes specifically on read alignments that can be affected by these genomes. Because the genomes used in this study are all haploid, we do not expect reads that overlap heterozygous variants to be significantly affected by the specific genome used.

We plotted the homozygous mapping error rates for the individual NA19238 (for each error type) in Figure 2e, and for all eight individuals (summed over all error types) in Figure 2f. Compared to Figure 2c-d, the homozygous error rates (Figures 2e-f) show a much steeper decrease when the Reference genome is replaced with the Pan-human consensus. Additionally, the heterozygous error rate is higher than the homozygous error rate and stays relatively constant across all genomes (Supplementary Figures S21-28). This supports the notion that consensus genomes significantly improve mapping accuracy of the reads that overlap homozygous variants, however, owing to their haploid nature, they cannot improve the alignment of the reads overlapping heterozygous loci.

## Mapping RNA-seq reads to unrelated consensus genomes outperforms the Reference

We next investigated the effects of mapping an individual’s RNA-seq reads to consensus genomes of different populations (Figure 3a) and to other personal haploid genomes (Figure 3c). We used the same reads, individuals, and genomes as previously discussed, and mapped all individuals to all genomes. The homozygous mapping error rate is calculated as before, and is shown in Figures 3b,d.

**Figure 3:**
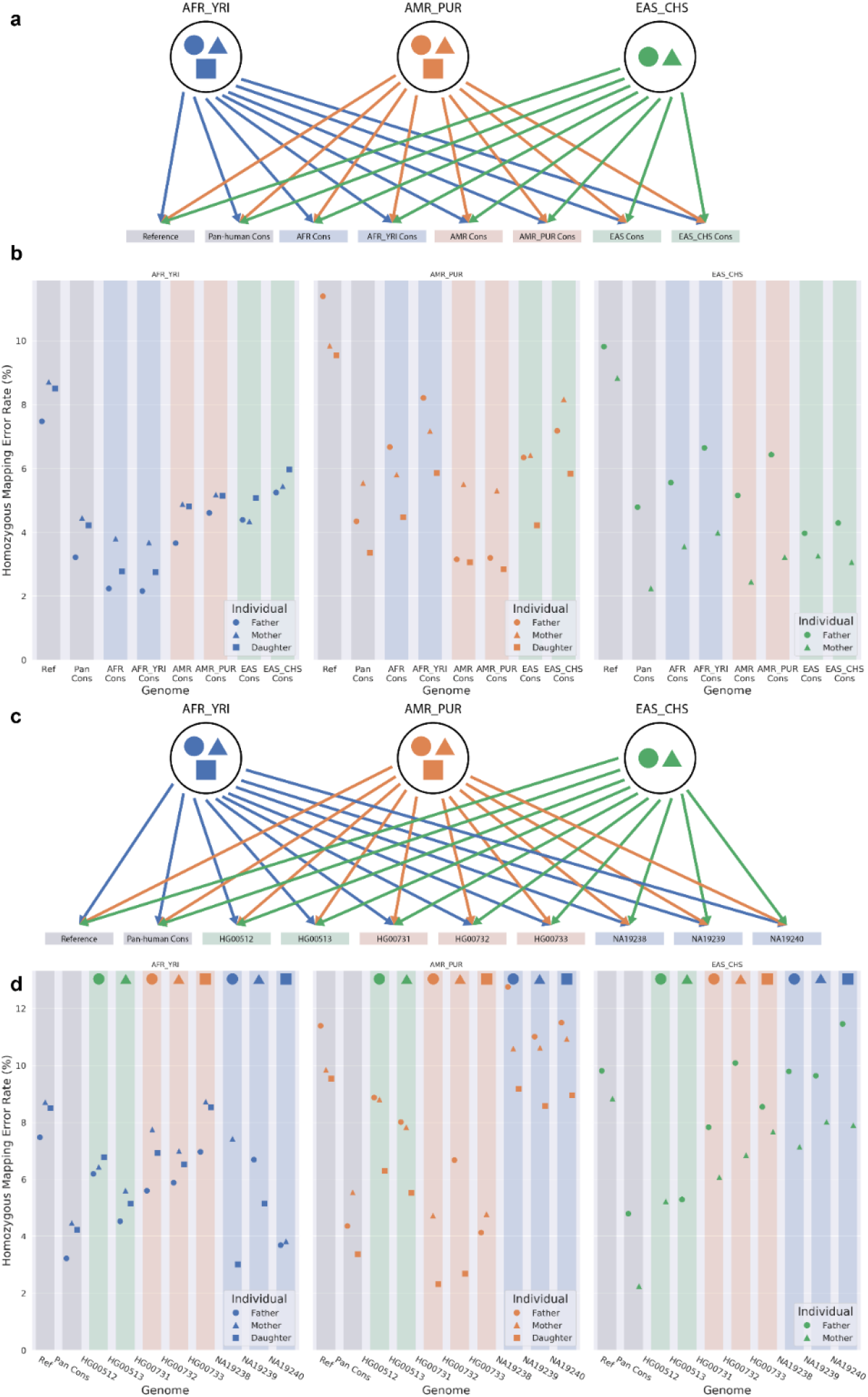
**a)** Each individual from each population is independently mapped to the Reference, Pan-human consensus, and all population and super-population consensus genomes. **b)** Homozygous mapping error rate when mapping to different consensus. The color of the marker indicates the population to which that individual belongs, while the shape of the marker identifies the individual within the trio. The color of the background rectangle indicates the population of the genome. **c)** Each individual from each population is independently mapped to the Reference, Pan-human consensus, and all personal haploid genomes. **d)** Homozygous mapping error rate when mapping to different personal haploid genomes. The color of the marker indicates the population to which that individual belongs, while the shape of the marker identifies the individual within the trio. The color of the background rectangle indicates the population of the genome. The shape at the top of each bar indicates to which individual in the trio that genome belongs.

As expected, Figure 3b shows that the unrelated consensus genomes perform worse than both related Population consensus and the Pan-human consensus, because each Population consensus contains many major alleles unique to that population. Interestingly, unrelated consensus genomes still perform better than the Reference. This is explained by the fact that the Reference contains a large number of minor alleles specific to the individuals who contributed to the Reference assembly. Conversely, the personal genomes of unrelated individuals are unlikely to share many MARs. This is illustrated in Figure 3d: the mapping error rate to personal genomes from different populations is higher than mapping to the Pan-human consensus and is comparable with mapping to the Reference. Notably, even mapping to the unrelated individual from the same population (Mother to Father and Father to Mother) does not improve the accuracy significantly. However, since the daughter in each trio will share many of her MARs with her parents, we see the error rates for mapping daughters’ RNA-seq reads to their parents’ genome (and vice versa) slightly better than mapping to the Pan-human consensus.

The results demonstrate that the Reference genome performs worse than any consensus genome, even consensuses from a different population. The accuracy of mapping to the Reference is comparable to mapping to unrelated personal genomes. On the other hand, the Pan-human consensus outperforms mapping to the unrelated individual genomes of the same or different population, and its performance is comparable with mapping to the genomes of related individuals (parent to child).

## Mapping error-causing variants are predominantly located in introns and UTRs

To investigate the genomic mechanisms underlying these mapping errors, we classified the genomic loci of the error-causing variants by overlapping error-causing reads with the GENCODE v29 GTF file. Interestingly, only a small proportion of the error-causing variants occur in the coding regions, while most are located in the intronic regions, followed by UTR and intergenic regions (Figure 4a). Because polyA+ RNA-seq reads should generally not map to introns, these errors are likely attributable to reads switching between being uniquely mapped and mapping to multiple locations (Unique to Multiple and Multiple to Unique error types). Interestingly, this corresponds with the previous observation that the largest sources of errors were the Unique to Multiple and Multiple to Unique error types.

**Figure 4:**
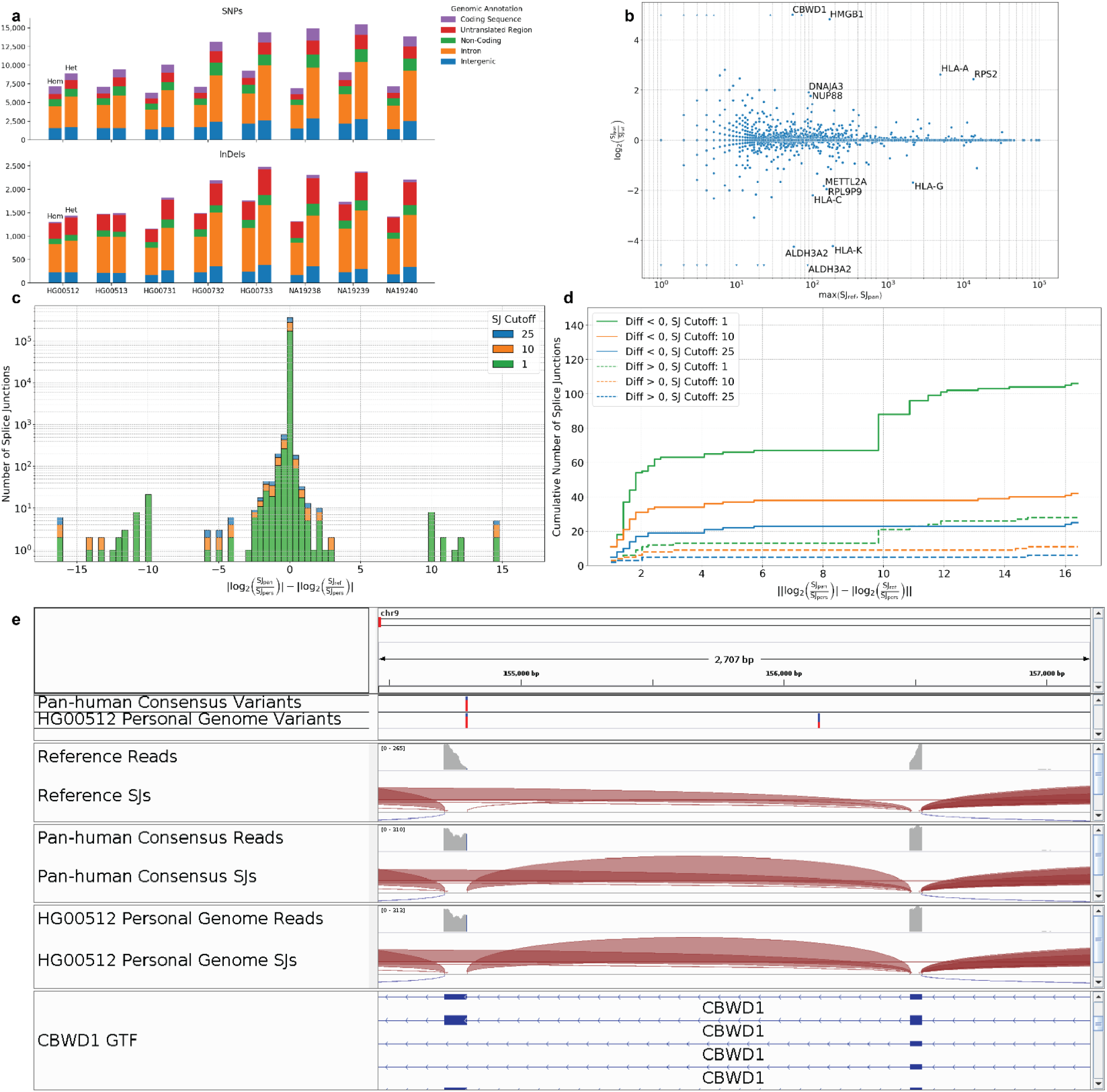
**a)** Counts of variants in the personal haploid genome that cause mapping errors in the Reference, classified by the genomic feature in which the variant is located. For each set of bars, the left bar shows the number of homozygous variants and the right bar shows the number of heterozygous variants. **b)** Log2 fold change between Pan-human consensus and Reference as a function of the max splice junction expression. Splice junctions with an absolute log fold change > 1.5 and a max expression value > 50 are labeled with the gene in which they fall. Triangles indicate an infinite log2 fold change (i.e. zero expression in one of the genomes). **C)** Difference between absolute values of Pan-human to Personal and Reference to Personal log-ratios. Different read count thresholds are represented by different colors. **D)** Cumulative distribution of the quantification error. Solid lines represent splice junctions which have larger quantification errors in the Reference than in the Pan-human genome; dashed lines represent the opposite cases. **e)** Read coverage and splice junction tracks for HG00512 reads aligned to the Reference, Pan-human consensus, and HG00512 personal genome. The region shown is part of the CBWD1 gene. The two Variants tracks show the location of a shared MAR that is present in the Pan-human consensus and the HG00512 personal genome.

## Consensus alleles generate large changes in splice junction expression

Here, we explore the effect of replacing the Reference with a consensus genome on splice junction expression. We define splice junction expression as the number of uniquely mapping reads which are spliced through the junction. Here we only consider annotated junctions, and define quantification error as the log2-ratio of the junction read counts in the Reference or Pan-human consensus to the junction read count in the personal genome (ground truth). Although the vast majority of splice junctions show very similar expression results for both genomes, there are many splice junctions with large quantification errors (Figures 4b-d). To reduce noise, we filtered the splice junctions with low expression in all three genomes at three counts thresholds of 1, 10, and 25 (Figure 4c-d). For all three thresholds, there were ~4-5 times as many splice junctions for which the quantification error in the Reference was higher than that in the Pan-human consensus (Figure 4d).

To illustrate the effect of consensus genomes on splice junction expression, we looked at a splice junction in the CBWD1 gene. This splice junction has very low expression in the Reference, but is highly expressed in the Pan-human consensus genome and the HG00512 personal genome. This disparity signifies a large error in the Reference with respect to the ground truth, which is mitigated by the Pan-human consensus. A genome browser snapshot of the region of the CBWD1 gene containing this splice junction is shown in Figure 4e, highlighting the effects that MARs can have on read mapping and on splice junction quantification. In this case, the Reference contains the minor allele, which prevents reads from mapping to the exon. However, both the Pan-human consensus and personal HG00512 genome contain the major allele, allowing the reads to be mapped to the exon. Because of this MAR, the isoforms containing this splice junction have erroneously low expression when reads are mapped to the Reference The Pan-human consensus rectifies the problem, predicting high expression of these isoforms that agrees with the ground truth of the personal genome mapping.

## Discussion

In any data analysis, often a first central question is how much variation to include. This might be accomplished by dimension reduction, quality control, feature selection, stratification, or other techniques. The human genome is no exception, and considering how best it should be summarized remains a crucial problem. Importantly, that problem may have a use-dependent solution: what is important for disease variant detection may not be important for RNA-seq alignment, and vice versa. The current Reference genome has had enormous utility, and before tearing down the infrastructure that has been built up to exploit it, it is important to consider alternatives carefully. Graph genome methods are one promising option, and they resolve the main deficiency in the reference: effectively incorporating all variation (or aspiring to). However, this comprehensiveness comes with its own host of issues, such as the lack of a simple coordinate system, difficulties with visualization, and significantly inflated computing requirements. The wide adoption of a graph-based reference genome will likely take a long time, given the history of switching from one version of the linear Reference to the next: GRCh38 was released in December 2013 (https://genome.ucsc.edu/FAQ/FAQreleases.html), and at the time of this writing, 7 years later, studies are still being published using GRCh37.

Although the full adoption of a graph genome may be several years in the future, the path there need not be a straight line. We may explore methods that partially improve on the current Reference, while imposing few of the costs of the graph methods. By progressively assessing the role of population variation (in essence, moving from low principal components to higher ones), we can develop intermediate forms moving from the current reference to more accurate reflections of population variation and, particularly, ones that still opt to summarize variability to some degree. The consensus genomes have substantial utility at the pan-human level, and then show a fall off past that point, suggesting that the Pan-human consensus can be considered a first step in the direction of adding population variation information to the Reference. Although consensus genomes are unable to comprehensively represent all human genotypic variation, they are still a desirable alternative to the Reference as they eliminate the millions of spurious minor alleles present in the current Reference genome, while maintaining a simple linear coordinate system.

In this study, we explored the advantages and limitations of using consensus genomes for RNA-seq mapping. We used read alignments to the haploid personal genome as a proxy for the ground truth to quantify the rate of erroneous alignments to the Reference genome, and compared it to the three levels of consensus: pan-human, super-population and population.

The overall mapping error rate caused by Reference shortcomings is quite small at only ~0.5-0.6% of all reads for the Reference genome, and further reduced to 0.3-0.4% for the consensus genomes, leaving relatively small room for further improvements (Figure 2d). However, for some analyses, such as allele-specific expression or de novo variant calling, the only reads of interest are those that overlap the variants. If we normalize the number of the erroneous reads by the number of reads that overlap the personal variants for each individual, we observe much higher corresponding error rates of ~8-10%, which decrease to ~2-4% when using a consensus genome.

The homozygous error rate (defined for reads that overlap only homozygous variants) is substantially decreased (by ~2-3 fold) when the Reference genome is replaced by the Pan-human consensus. Surprisingly, using the Super-population or Population consensuses does not result in further improvement of the mapping accuracy, which indicates that the Pan-human consensus captures the majority of population variation information that can be captured in a linear haploid genome. Using the Super-population or Population consensus genomes may not be worth the loss of generality: for instance, it will severely complicate interpopulation comparisons owing to the lack of a common coordinate system.

These mapping results call into question the time and resources that are being spent on constructing consensus genomes for particular populations (Cho et al. 2016; Fakhro et al. 2016; Higasa et al. 2016; Sherman et al. 2019; Takayama et al. 2019). Intuitively, one would expect that more specific consensus genomes would increase the mapping accuracy for the populations that they represent. However, our results indicate that a universal Pan-human consensus genome is sufficient to attain the best possible accuracy that can be achieved with a haploid reference, and the expensive efforts to construct more population-specific references are likely futile for improving accuracy of RNA-seq analyses.

On the other hand, the heterozygous error rate (for reads that overlap heterozygous variants) is not significantly reduced by replacing the Reference with a consensus of any population level. This is not surprising given that the haploid genome can only include one of the alleles of a heterozygous locus, and hence cannot truthfully represent it. Graph genomes or other non-linear reference representations will be required to reduce error rates for heterozygous loci.

Although there is still work to be done on improving the Reference genome, the Pan-human consensus already offers noticeable improvements in downstream analyses, as indicated by the difference in splice junction expression quantification. We demonstrated that the accuracy of the splice junction quantification is significantly improved by switching from the Reference to the Pan-human consensus. These improvements imply important consequences in functional analyses such as alternative splicing, transcript abundance quantification and differential isoform usage. Splice junction differences are subtle, but the 5-fold difference in the number of splice junctions with higher quantification error in the Reference than in the Pan-human consensus demonstrates that the Pan-human consensus offers important improvements over the Reference. Results from a similar analysis of gene isoform expression (Supplementary Information) provide additional support for this claim.

The Pan-human consensus appears to be a strict improvement over the current Reference with minimal costs, and thus we propose replacing the current Reference with the Pan-human consensus. Besides the question of absolute utility, we also advocate using consensus genomes as a mechanism to develop practices to improve genome representation more generally. Recent years have seen genomics pipelines using the Reference become entrenched, to varying degrees, by researchers unwilling to upgrade. Because the consensus genome requires very minor changes in pipelines, it can be used as a straightforward, first-order approximation to assess and explore the sensitivity of specific genomic analyses to genome variation. For instance, the benefits of the consensus genome for RNA-seq mapping can be explored via the STAR-consensus pipeline, which aligns reads to the consensus genome and then transforms the coordinates to the Reference genome coordinates, thus eliminating the need for changes in the downstream processing. By incorporating consensus genomes, we envision not only improvements in both the absolute performance of diverse research projects, but also a greater understanding of the dependencies in those methods, thus setting the stage for a more flexible and robust future for genomics.

## Methods

### Calculating consensus alleles

We calculated the consensus allele for each variant on a per-haplotype basis: the number of occurrences of each allele was counted, and the most common allele was selected. For the Pan-human consensus, the alleles were counted across all individuals. For each Super-population and Population consensus, the alleles were counted across all of the individuals within that group. This counting was performed in Python by ConsDB, by reading through each VCF file one line at a time and parsing the genotype for each individual in the group for which the consensus is being constructed.

### Genome generation and read mapping

All genomes generation and read mapping was done with STAR v2.7.7a (Dobin et al. 2013). We used GRCh38 (Schneider et al. 2016) as the reference FASTA file and GENCODE v29 (Frankish et al. 2019) as the reference GTF file. We masked the PAR regions on the Y chromosome in order to avoid any sex-based differences in mapping. For the generation of consensus and personal haploid genomes, we used the `--genomeTransformType Haploid` option and the `--genomeTransformVCF` option with the appropriate VCF file. For the read mapping, we used the `--genomeTransformOutput SAM SJ` and the `--quantMode GeneCounts TranscriptomeSAM` options. We also used the `--outSAMreadID Number` option in order to more easily keep track of reads in the analysis steps. Other than these options, we used the default STAR parameters.

### Mapping error calculations

Before calculating the mapping error, we made a number of preparations. First, we used awk to construct a VCF file for each individual that contained only the variants and that individual’s phased genotype. Next, we used these full VCFs to partition the variants for each consensus genome for each individual into four separate VCF files: one for homozygous SNPs, one for heterozygous SNPs, one for homozygous indels, and one for heterozygous indels. These four split VCFs needed to be generated for each individual, including individuals from within the same population, because variants may be homozygous in one individual but heterozygous in a different individual.

For each individual, their filtered alignments for the Reference, Pan-human consensus, Super-population consensus, and Population consensus were compared to the filtered alignment for their personal haploid genome using an awk script. We compared the genomes on a per-read basis, checking for differences in mapping position and number of mapped loci. To determine what types of variants each read overlapped, we overlapped the filtered BAM files with each of the four split VCF files using bedtools, for each genome and each individual. We compared the read IDs from this overlap with the read IDs obtained from the genome mapping comparisons using grep, in order to find error-causing variants.

The final steps of read counting and plotting were done using a Python script. For each individual, we summed the read counts for each combination of error type and homozygous/heterozygous variants across all four genomes being analyzed. The two normalization constants used for these figures were the total number of mapped reads for each individual, and the total number of reads for each individual that overlapped personal homozygous variants. The total mapped read numbers were extracted from the STAR Log.final.out file. The counts of reads overlapping personal homozygous variants were found by counting the number of reads present in the previously found overlap files for reads overlapping homozygous variants in the personal haploid genome.

### Finding error-causing variant locations

To find the genomic annotations of error-causing variants, we first found the error-causing variants as described previously. We next used bedtools to intersect these variants with the GENCODE v29 (Frankish et al. 2019) GTF file and find all genomic annotations that each variant overlaps. Because certain genomic annotations always fall within other genomic annotations (e.g. an exon will necessarily be located within a gene), a given variant is likely to have multiple genomic annotations that it overlaps. We used a Python script to determine the most specific genomic annotation overlapped by each variant and to count the number of variants falling within each type of genomic annotation.

### Splice junction expression calculations

The splice junction expression values used in our calculations were generated during the previously described read mapping section. Specifically, we used the number of uniquely mapped reads crossing the splice junction, which is given in the 7th column of the SJ.out.tab file generated by STAR. To calculate the quantification error for each genome, we used custom Python scripts. The log2 fold change values shown in Figure 4b were plotted without normalization, and the splice junctions for which the Reference had no unique reads crossing the splice junction were represented as arrows. Splice junctions for which both the Pan-human consensus and the Reference had 0 expression were excluded from this plot. Additionally, these log2 fold change values were thresholded to +/− 5. Splice junctions with an absolute log fold change > 1.5 and a max expression value > 50 were labeled with the gene in which they fall. In Figures 4c-d, all read count values were normalized by an addition of 0.001 in order to prevent infinite log2 fold change values.

### Selection of splice junction of interest

We selected the splice junction through a manual inspection process. We searched for a splice junction with a large absolute log2 fold change between the Reference and the Pan-human consensus, in order to find an example that would highlight the differences in splice junction expression between the two genomes. We also required that the splice junction fall within a protein coding gene.

### Code Availability

The ConsDB package is available on GitHub at https://github.com/kaminow/consdb. STAR-consensus is available at https://github.com/alexdobin/star. Scripts to re-produce the analysis in this study are available at https://github.com/kaminow/ConsDB_analysis.

