## Supplementary Information for "Virtue as the mean: Pan-human consensus genome significantly improves the accuracy of RNA-seq analyses"

### Consensus allele changes in isoform expression

We used RSEM (Li and Dewey 2011) to quantify transcript expression (TPM) for reads mapped to the Reference and to the Pan-human consensus, again using the haploid personal genome as the ground truth. In this analysis, we excluded insertions and deletions in order to avoid any discrepancies between the internal transformed genome generated by STAR and the genome used by RSEM to generate its genomic indices. The quantification error is defined as the log2-ratio of the TPM in the Reference or Pan-human consensus to the TPM in the personal (ground truth) genome. Although the vast majority of transcripts show very similar expression results for both genomes, there are many transcripts with large quantification errors (Supplementary Figure S29a-c). To reduce noise, we filtered the transcripts with low expression in all three genomes at three TPM thresholds of 0.2, 1 and 5 (Supplementary Figure S29b-c). For all three thresholds, there were ~6 times as many transcripts for which the quantification error in the Reference was higher than that in the Pan-human consensus (Supplementary Figure S29c).

To illustrate the effect of consensus genomes on the transcript quantification, we looked at a transcript of the ALDH3A2 gene, which has zero expression in the personal genome and the Pan-human consensus genome (no error), but at the same time exhibits non-zero expression in the Reference (~2 TPM), which signifies a large error with respect to the ground truth. A genome browser snapshot of selected regions of this gene is shown in Supplementary Figure S29d, highlighting the effects that MARs can have on read mapping and on transcript expression prediction. In this case, the Reference contains the minor allele, which causes reads to map to the short exon, and hence the isoform (ENST00000582991.5) containing this exon has non-zero expression. At the same time, both the Pan-human consensus and personal HG00512 genome contain the major allele, which prompts the read alignments to skip the short exon, resulting in zero expression for the ENST00000582991.5 isoform.

### Isoform expression calculations

The genome generation and mapping procedures for the transcript expression calculations were similar to the procedures for the mapping error section, however there were some key differences. The main difference in the genome generation step was the exclusion of insertions and deletions from the Pan-human consensus and HG00512 personal haploid genome. We excluded insertions and deletions for this analysis in order to avoid any discrepancies between the internal transformed genome generated by STAR and the genome used by RSEM to generate its genomic indices. The VCF files used to generate the Pan-human consensus and the HG00512 personal haploid genome were generated by using an awk script to remove insertions and deletions from the VCF files that were used to generate the genomes for the mapping error section. We generated the STAR genomes as previously described, using the SNP-only VCF files. We generated the RSEM genome indices using transformed genome FASTA files, which were made using bcftools to incorporate the SNPs in the VCF files into the PAR-masked genome.

The mapping steps for this analysis were the same as previously described, with the addition of the `--quantMode GeneCounts TranscriptomeSAM` option to force STAR to export transcriptomic alignments in addition to the standard genomic alignments. We ran RSEM using these STAR-generated transcriptomic alignments as the input.

We used Python scripts to calculate the transcript expression error for each genome, using the TPM values from the RSEM predictions. The log2 fold change values shown in Supplementary Figure S29a were plotted without normalization, and the transcripts for which either the Pan-human consensus or the Reference had an estimated TPM of 0 were represented as arrows. Transcripts for which the personal genome had an estimated TPM of 0 were excluded because these transcripts would, by definition, have an infinite log2 fold change for both the Pan-human consensus and the Reference. Additionally, transcripts for which both the Pan-human consensus and the Reference had an estimated TPM of 0 were excluded in this plot. In Supplementary Figure S29b-c, all TPM values were normalized by an addition of 0.001 in order to prevent infinite log2 fold change values.

### Selection of transcript of interest

We selected transcript ALDH3A2-222 through a manual inspection process. We searched for a transcript with significant differential expression between the Reference and the Pan-human consensus, in order to find an example that would highlight the differences in transcript expression calculation between the two genomes. We used DESeq2 to determine statistical significance. However, DESeq2 requires replicates in its inputs, and the HGSV reads only contained one sample. In order to utilize DESeq2 with this data, we randomly split the transcriptomic alignment of each genome into two separate BAM files, ensuring that all alignments for a given read were grouped in the same file. This splitting was performed using a Python script. We then ran RSEM on these split BAM files, and used the expected_count column of the RSEM output as the input for DESeq2. Because this column can contain non-integer values, all expected_count values were rounded down to the previous integer. For our selection, we only considered transcripts with a DESeq2 adjusted p-value $< 0.05$. In addition, we also required that the gene to which the transcript belongs was protein coding.
