## Supplementary Figures for "Virtue as the mean: Pan-human consensus genome significantly improves the accuracy of RNA-seq analyses"

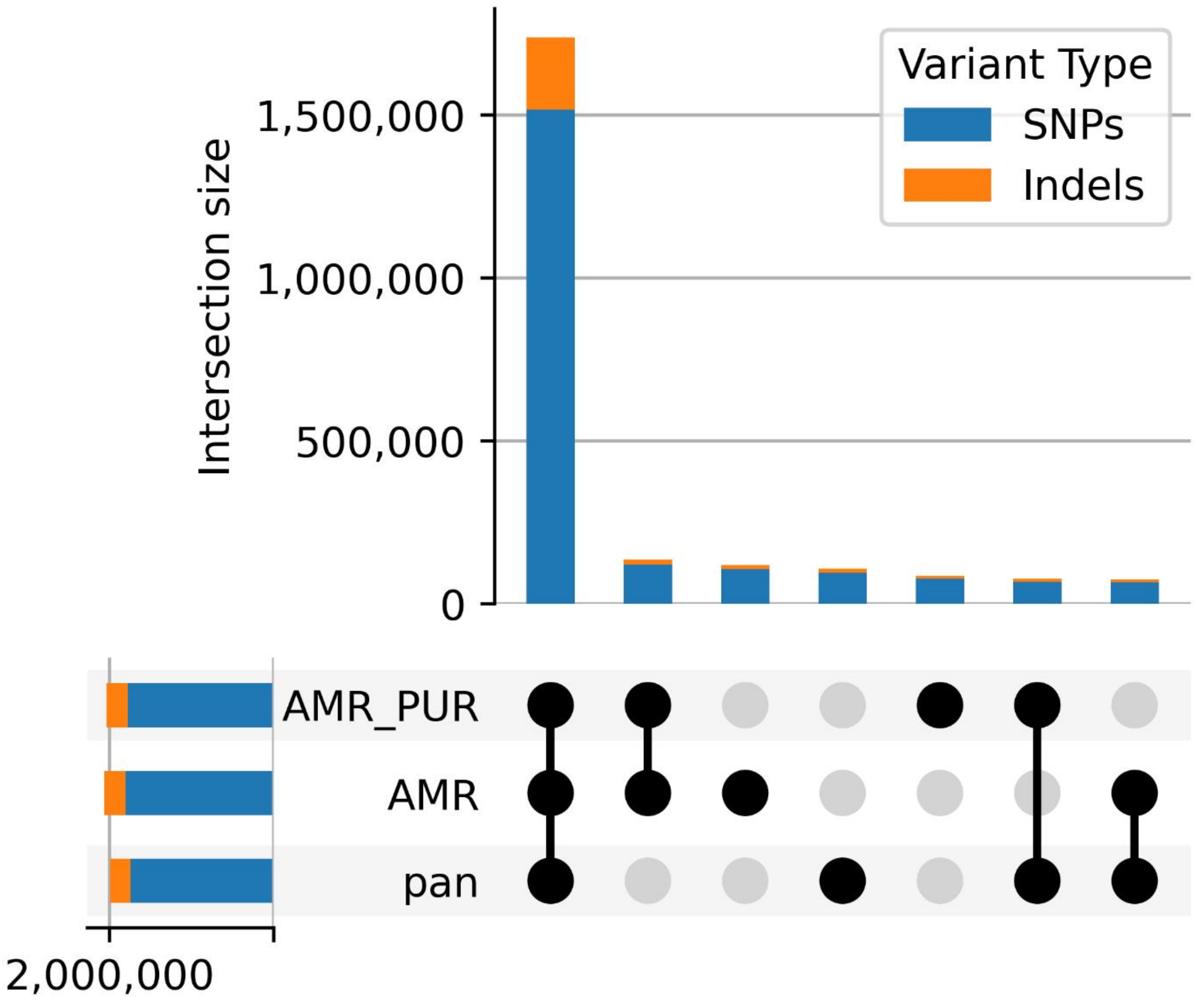

**Supplementary Figure S1:** Number of SNPs and indels shared between different combinations of the Pan-human, Super-population, and Population consensus genomes for the Admixed American population.

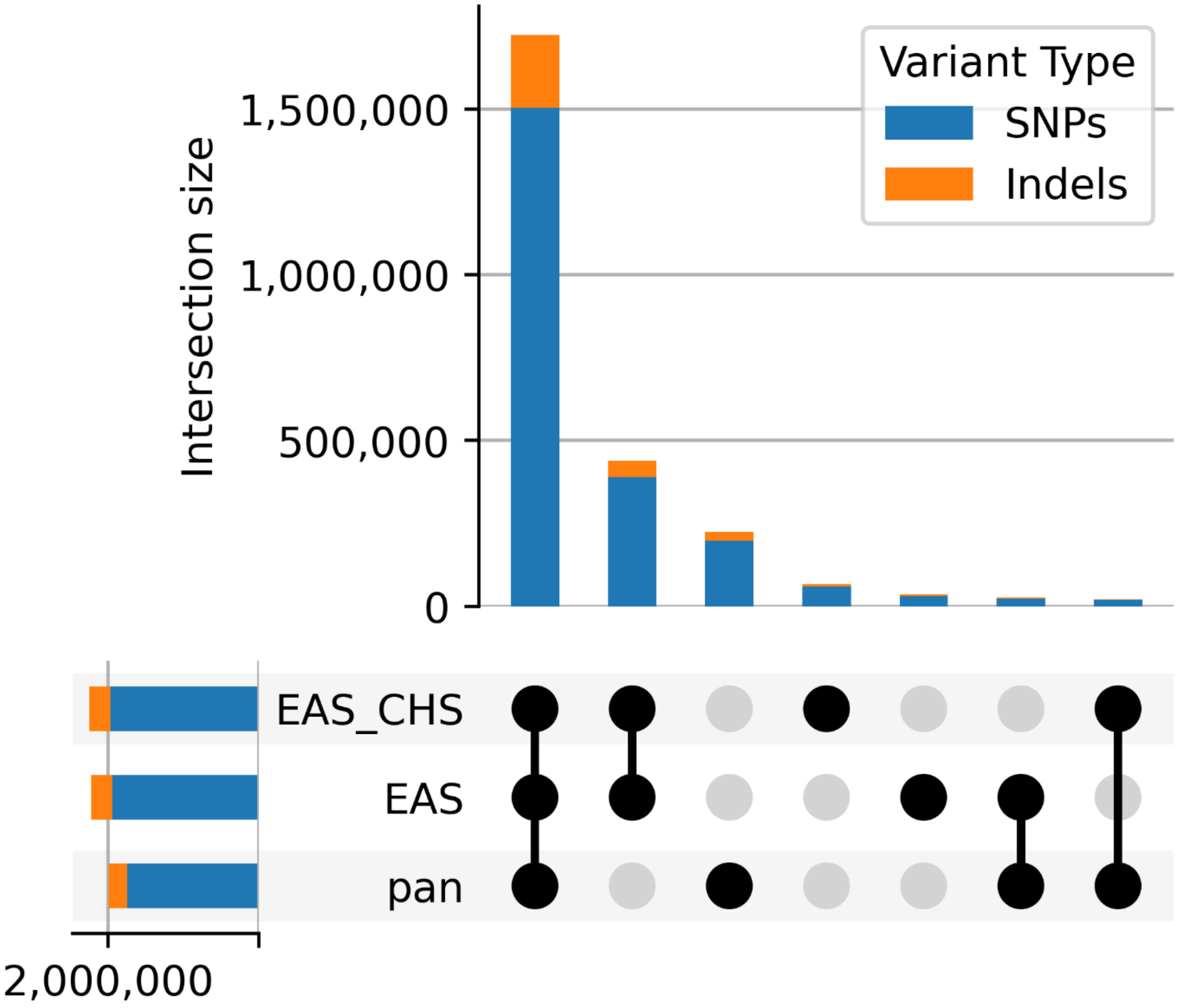

**Supplementary Figure S2:** Number of SNPs and indels shared between different combinations of the Pan-human, Super-population, and Population consensus genomes for the East Asian population.

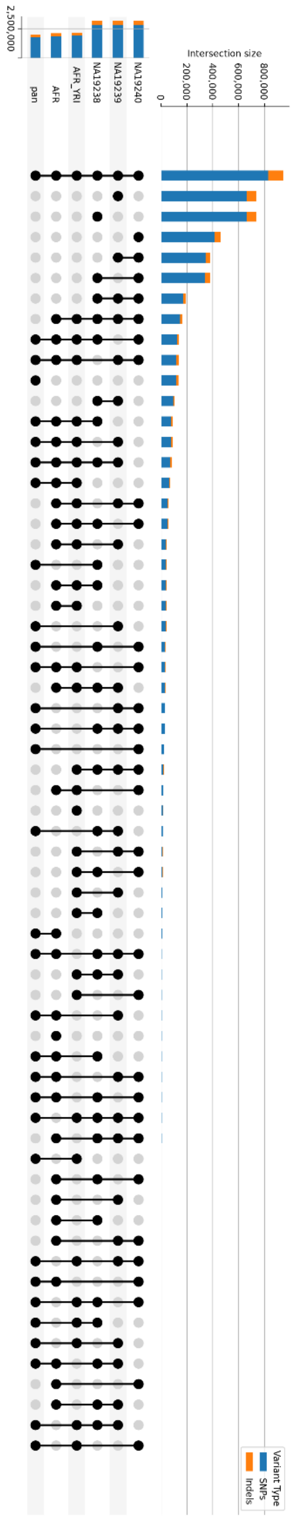

**Supplementary Figure S3:** Number of SNPs and indels shared between different combinations of the Pan-human, Super-population, and Population consensus genomes, and individual haploid personal genomes for the African population.

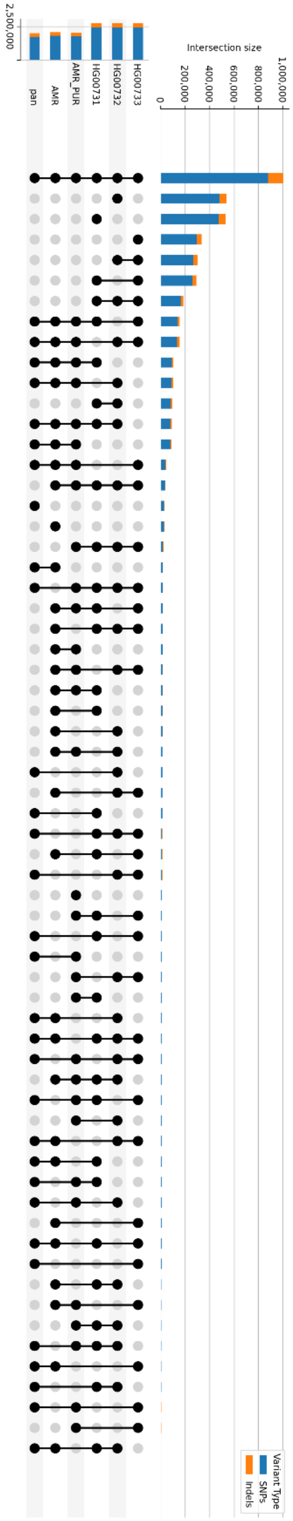

**Supplementary Figure S4:** Number of SNPs and indels shared between different combinations of the Pan-human, Super-population, and Population consensus genomes, and individual haploid personal genomes for the Admixed American population.

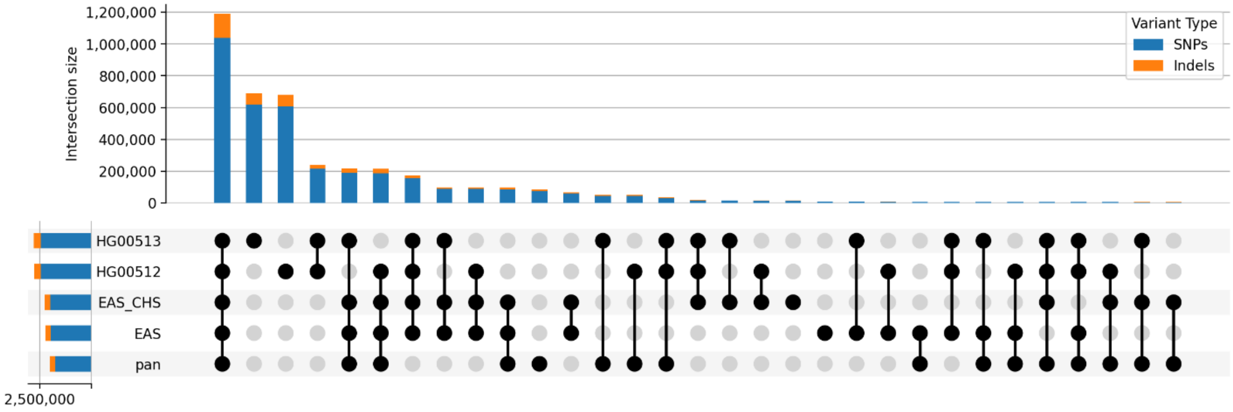

**Supplementary Figure S5:** Number of SNPs and indels shared between different combinations of the Pan-human, Super-population, and Population consensus genomes, and individual haploid personal genomes for the East Asian population.

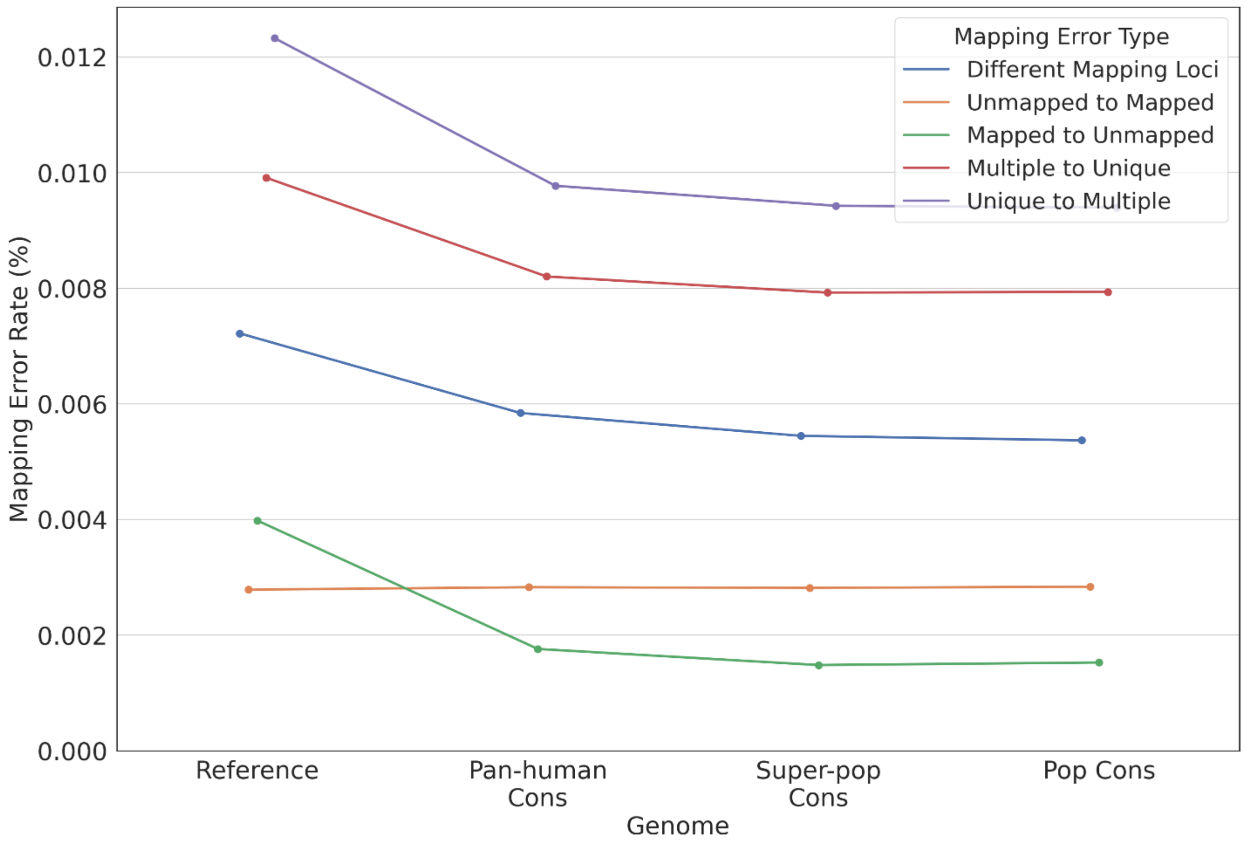

**Supplementary Figure S6:** Overall mapping error rate of reads overlapping insertions or deletions for each error type for individual NA19238.

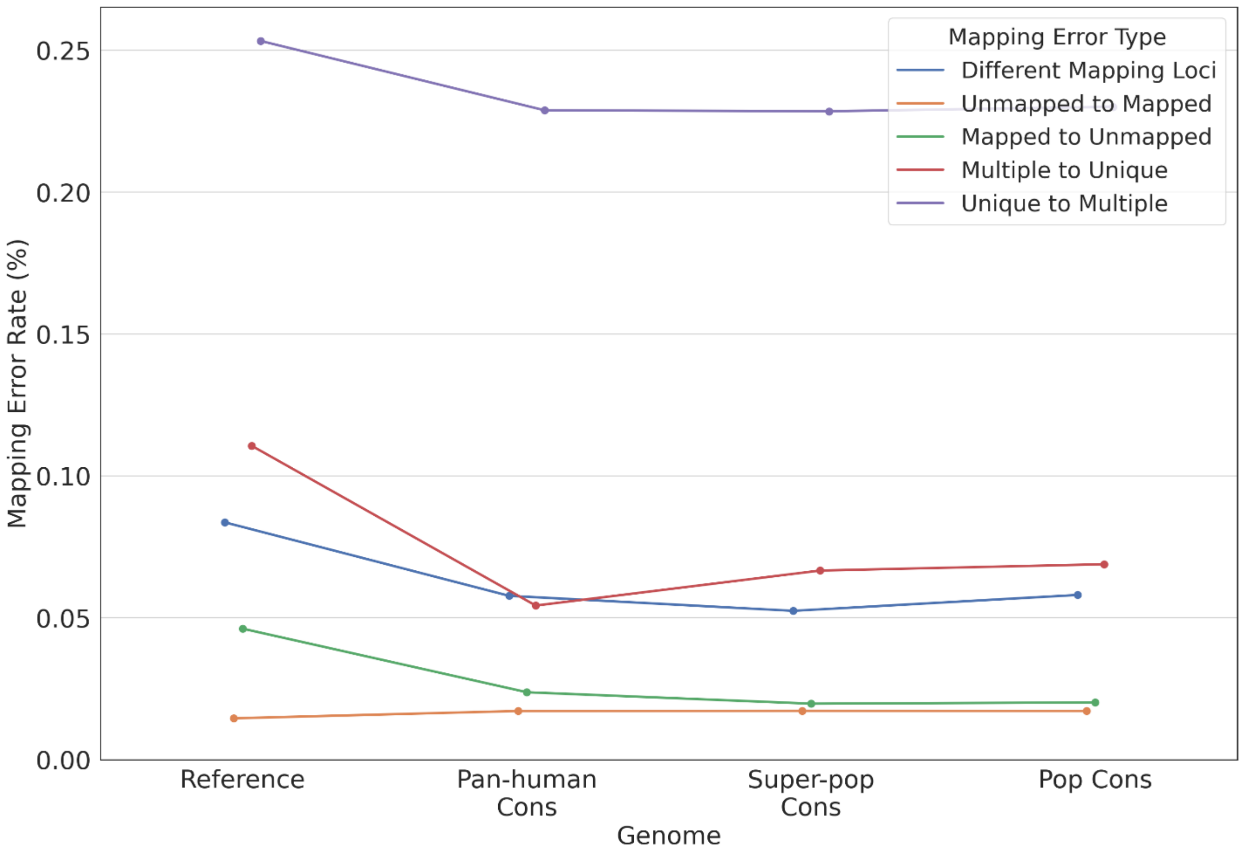

**Supplementary Figure S7:** Overall mapping error rate for each error type for individual HG00512.

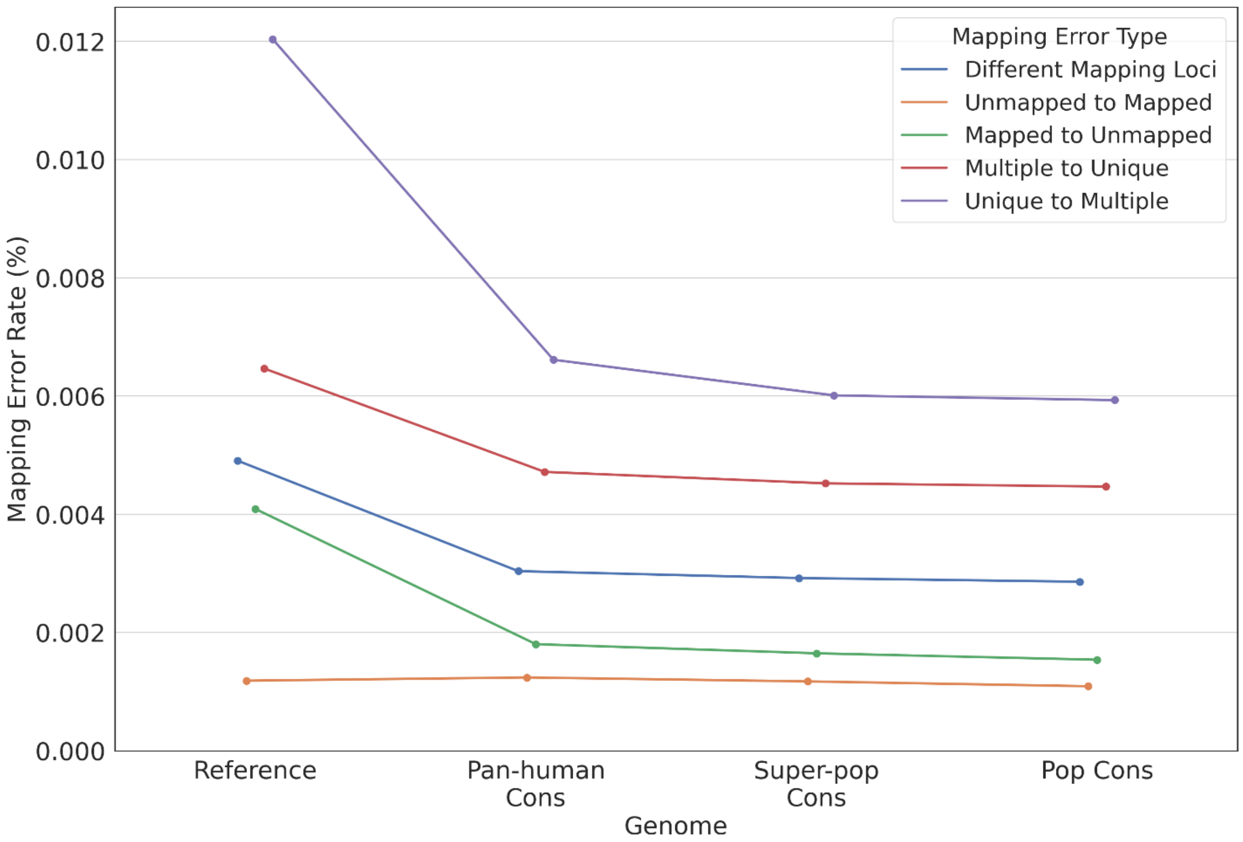

**Supplementary Figure S8:** Overall mapping error rate of reads overlapping insertions or deletions for each error type for individual HG00512.

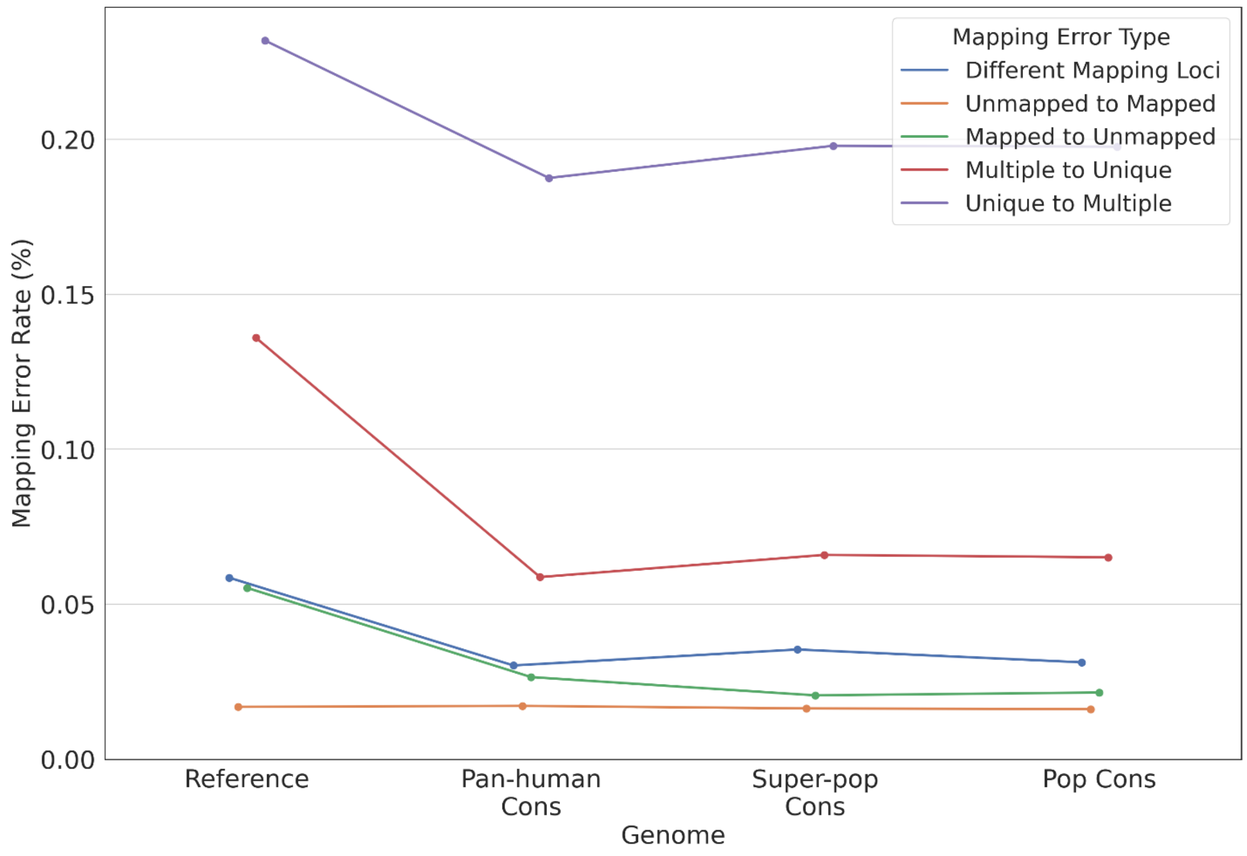

**Supplementary Figure S9:** Overall mapping error rate for each error type for individual HG00513.

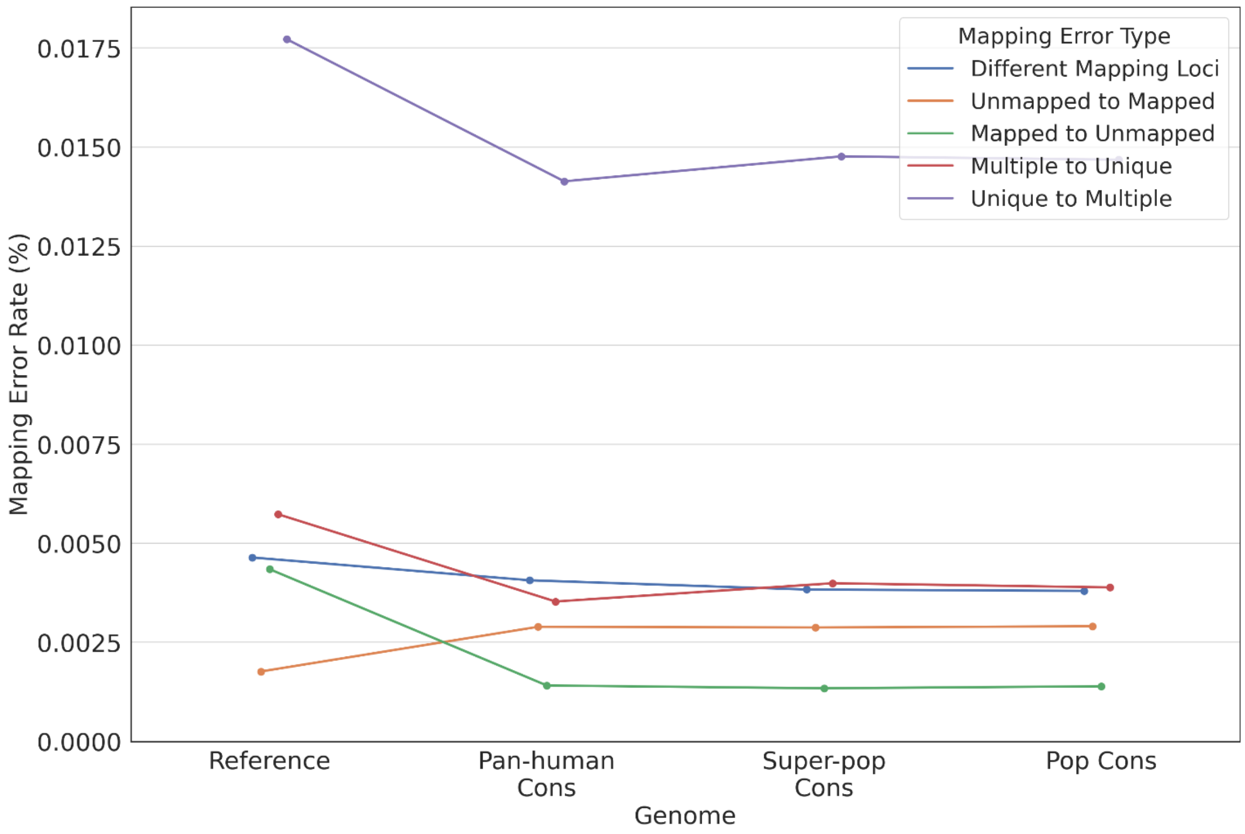

**Supplementary Figure S10:** Overall mapping error rate of reads overlapping insertions or deletions for each error type for individual HG00513.

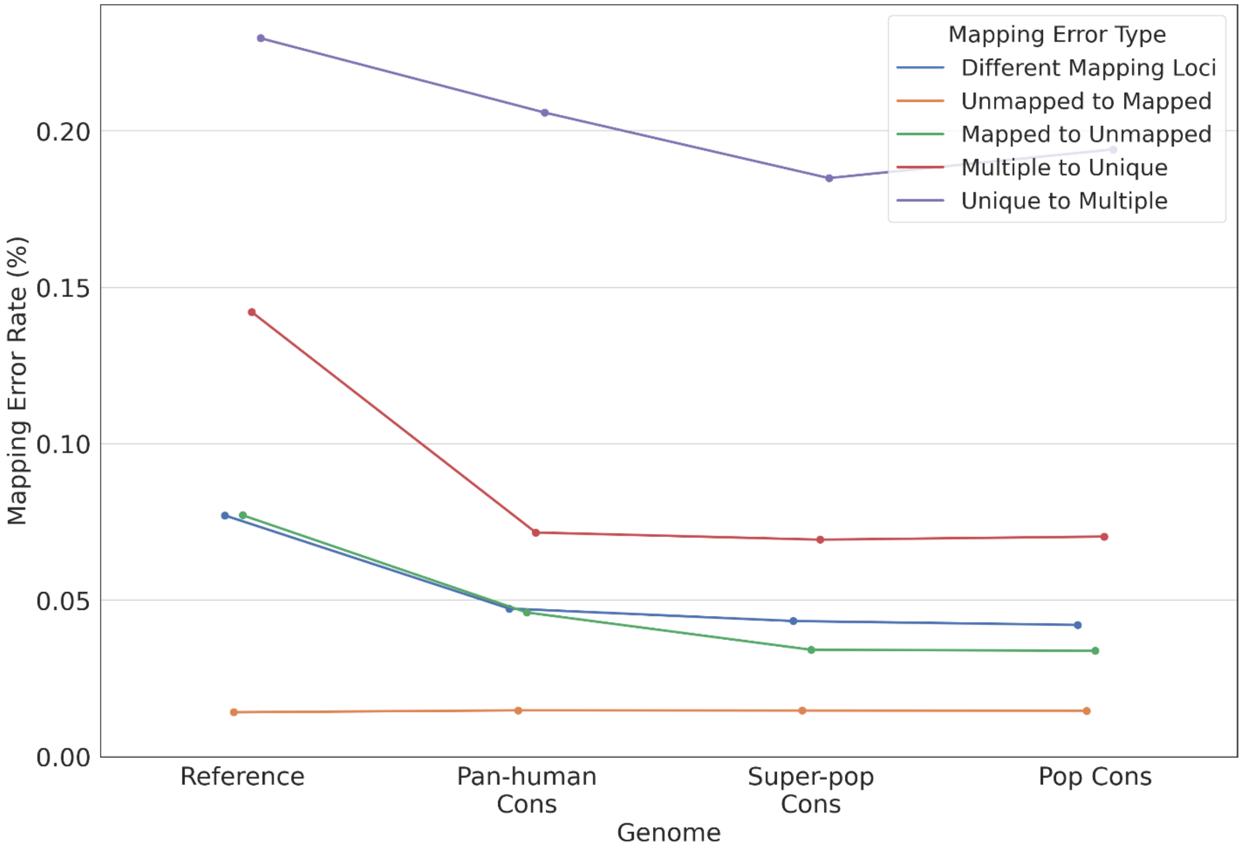

**Supplementary Figure S11:** Overall mapping error rate for each error type for individual HG00731.

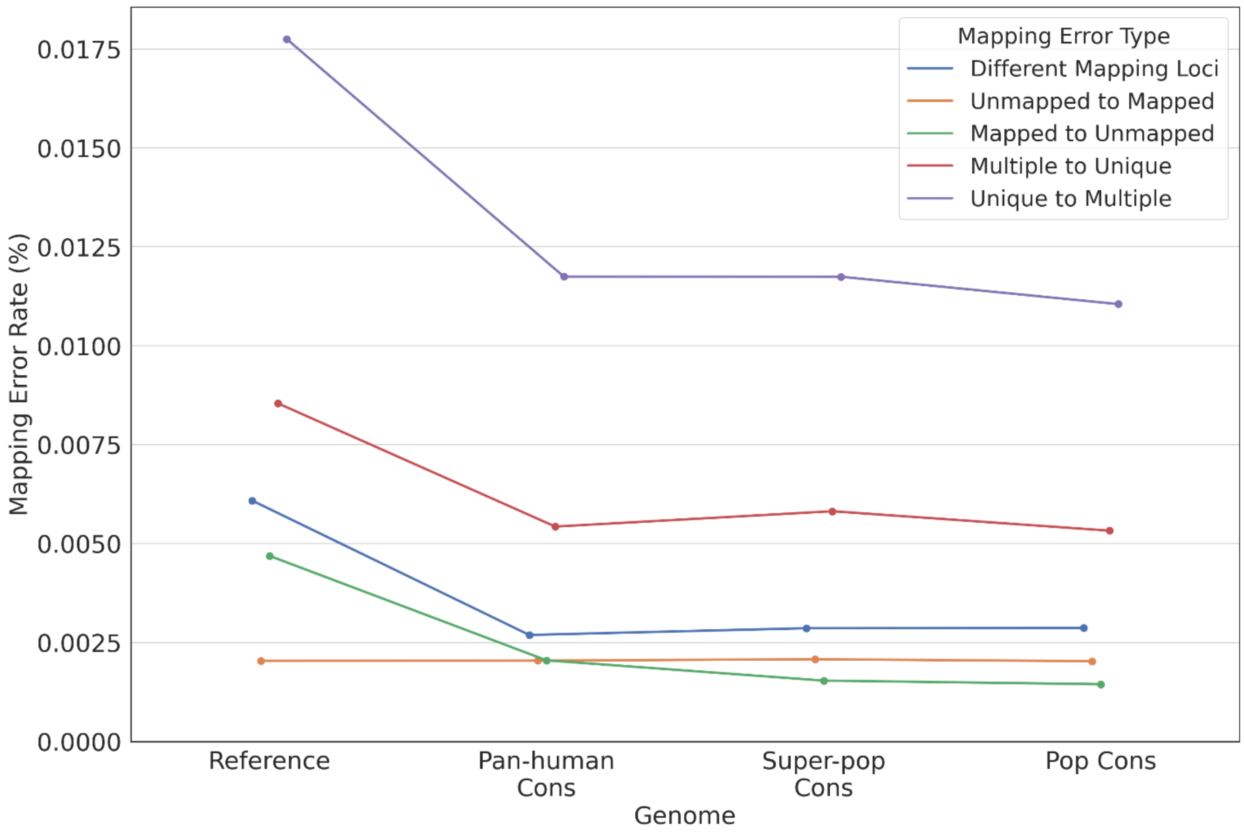

**Supplementary Figure S12:** Overall mapping error rate of reads overlapping insertions or deletions for each error type for individual HG00731.

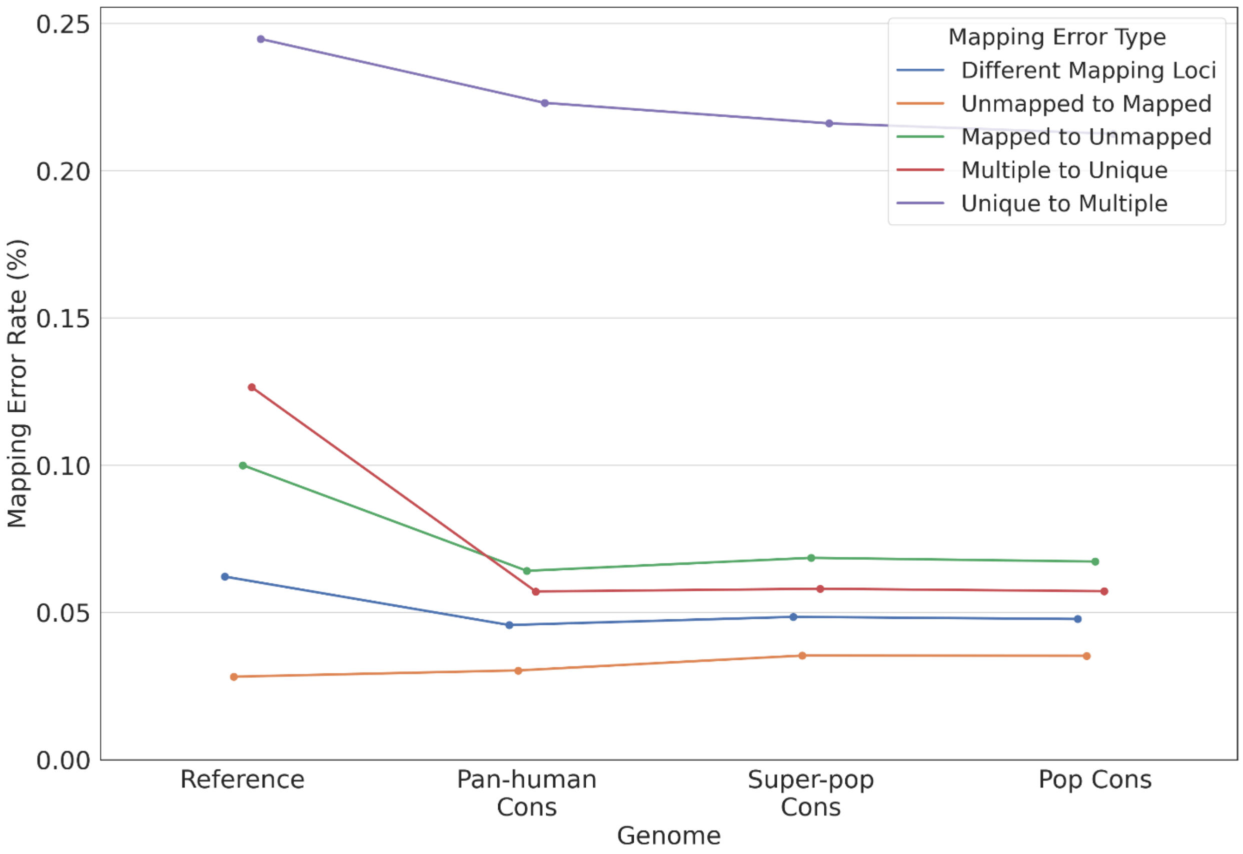

**Supplementary Figure S13:** Overall mapping error rate for each error type for individual HG00732.

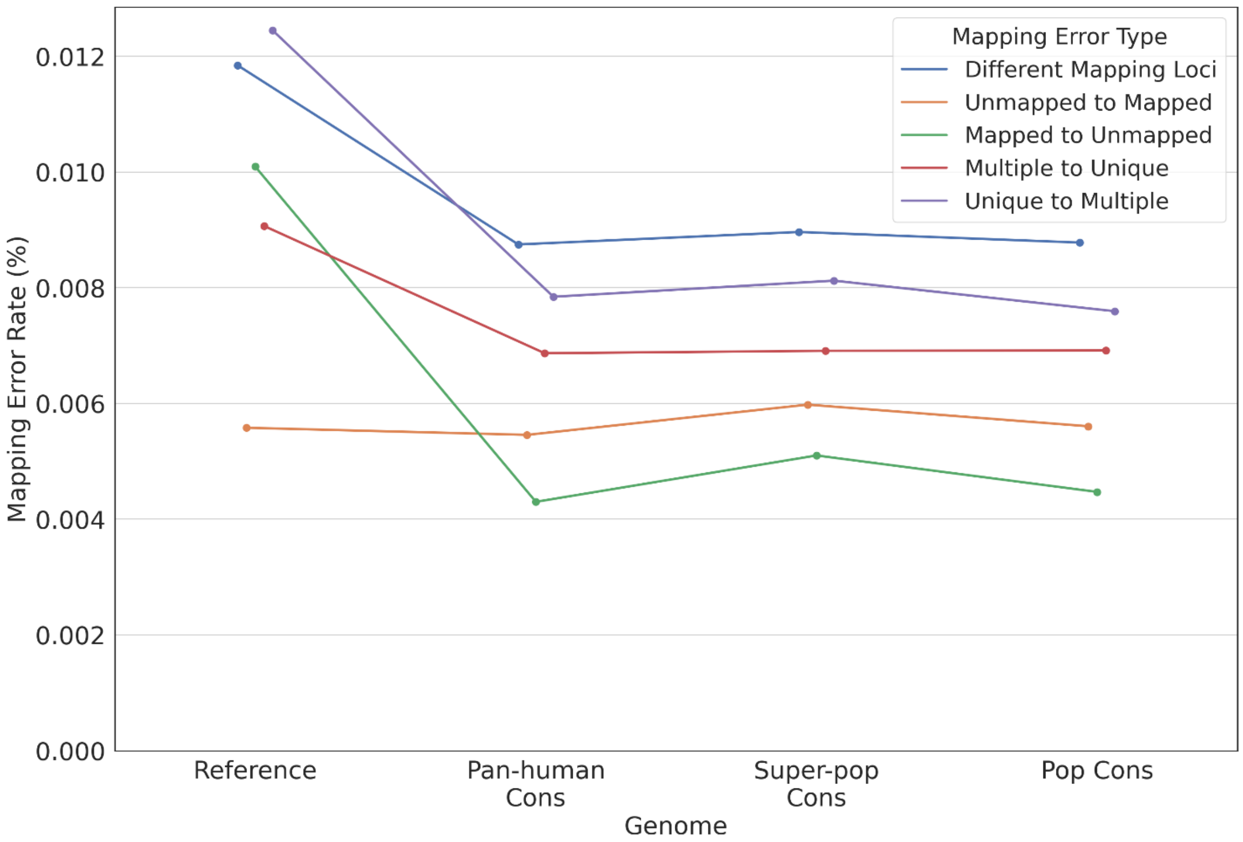

**Supplementary Figure S14:** Overall mapping error rate of reads overlapping insertions or deletions for each error type for individual HG00732.

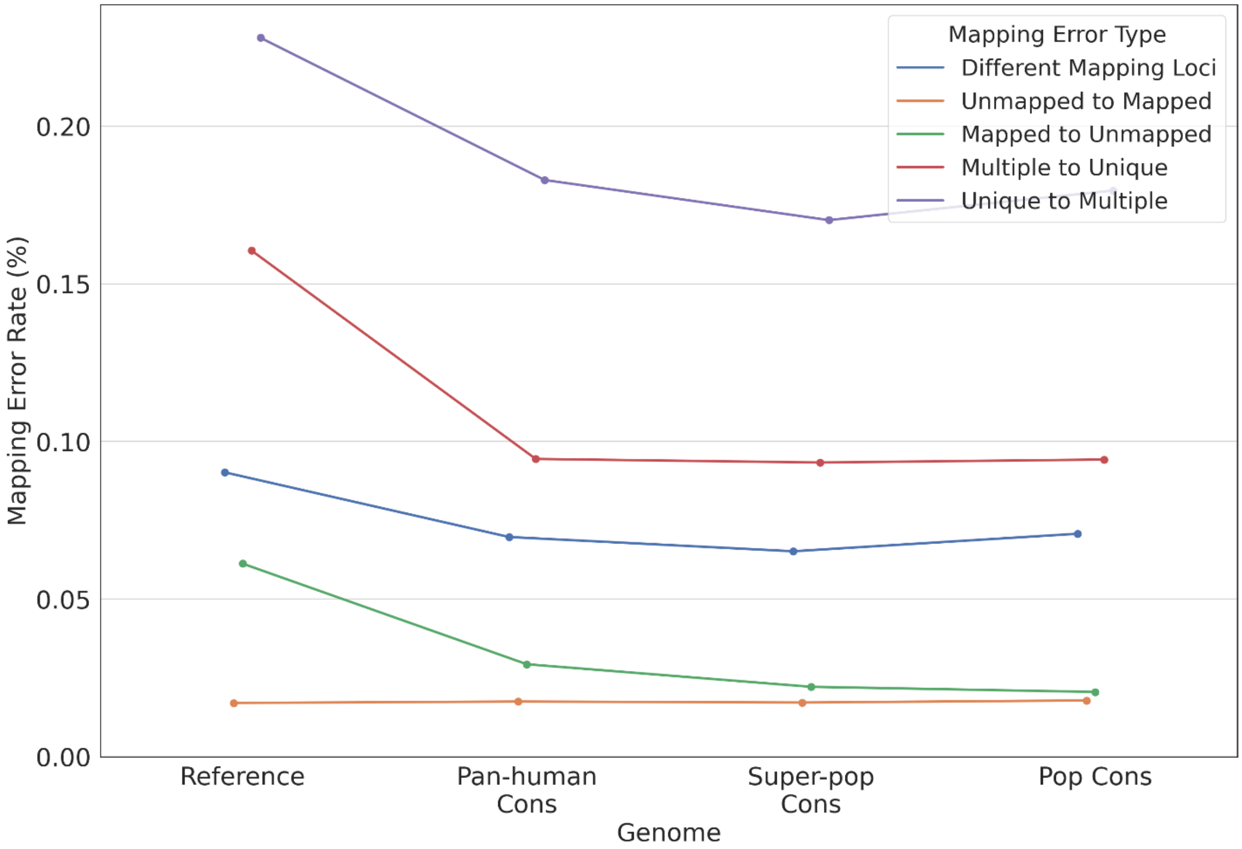

**Supplementary Figure S15:** Overall mapping error rate for each error type for individual HG00733.

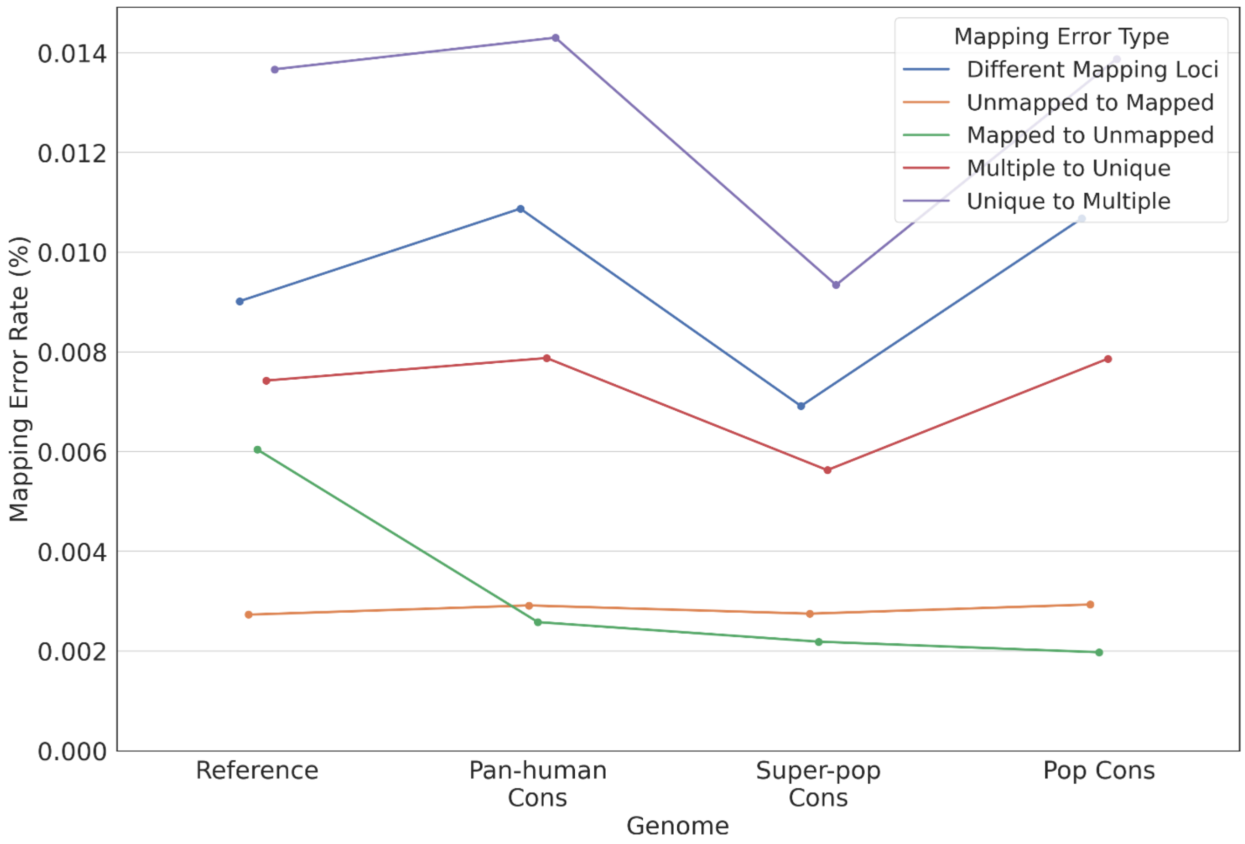

**Supplementary Figure S16:** Overall mapping error rate of reads overlapping insertions or deletions for each error type for individual HG00733.

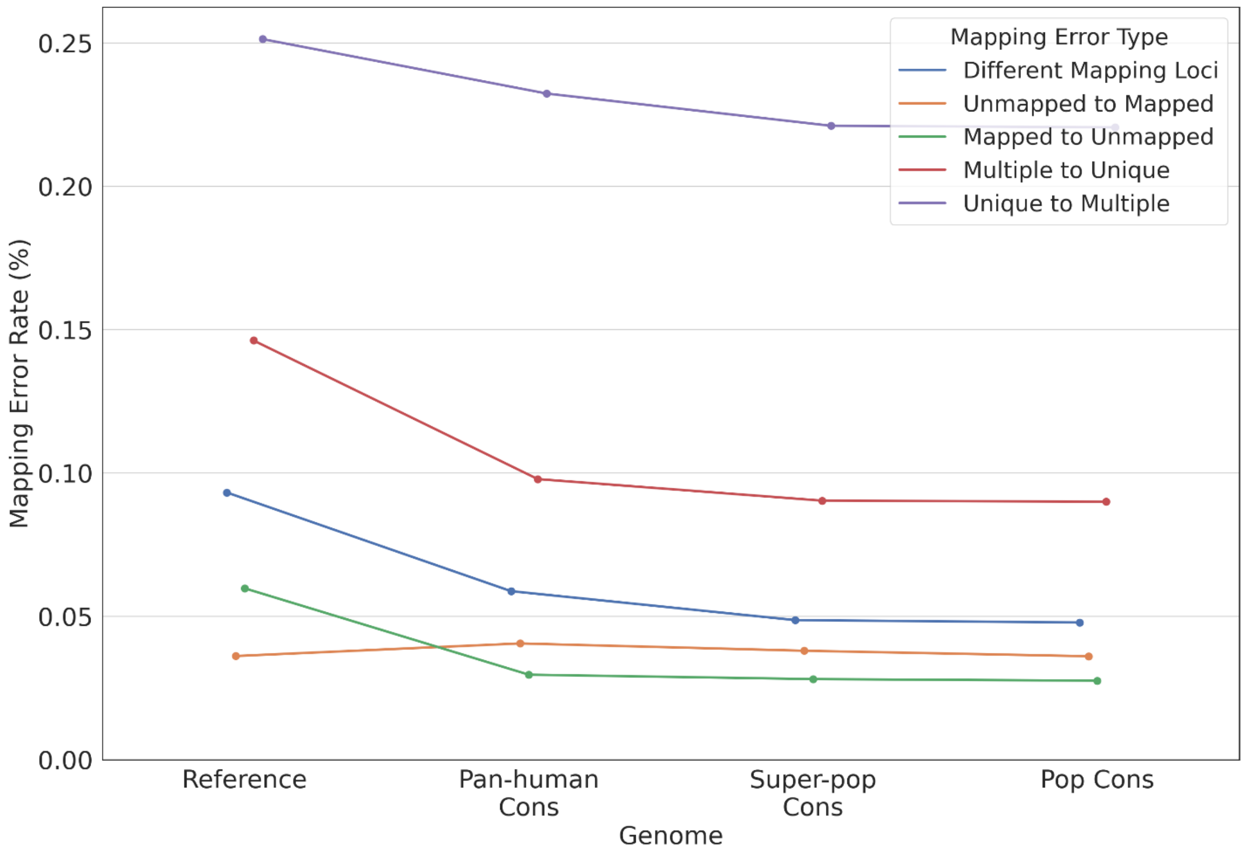

**Supplementary Figure S17:** Overall mapping error rate for each error type for individual NA19239.

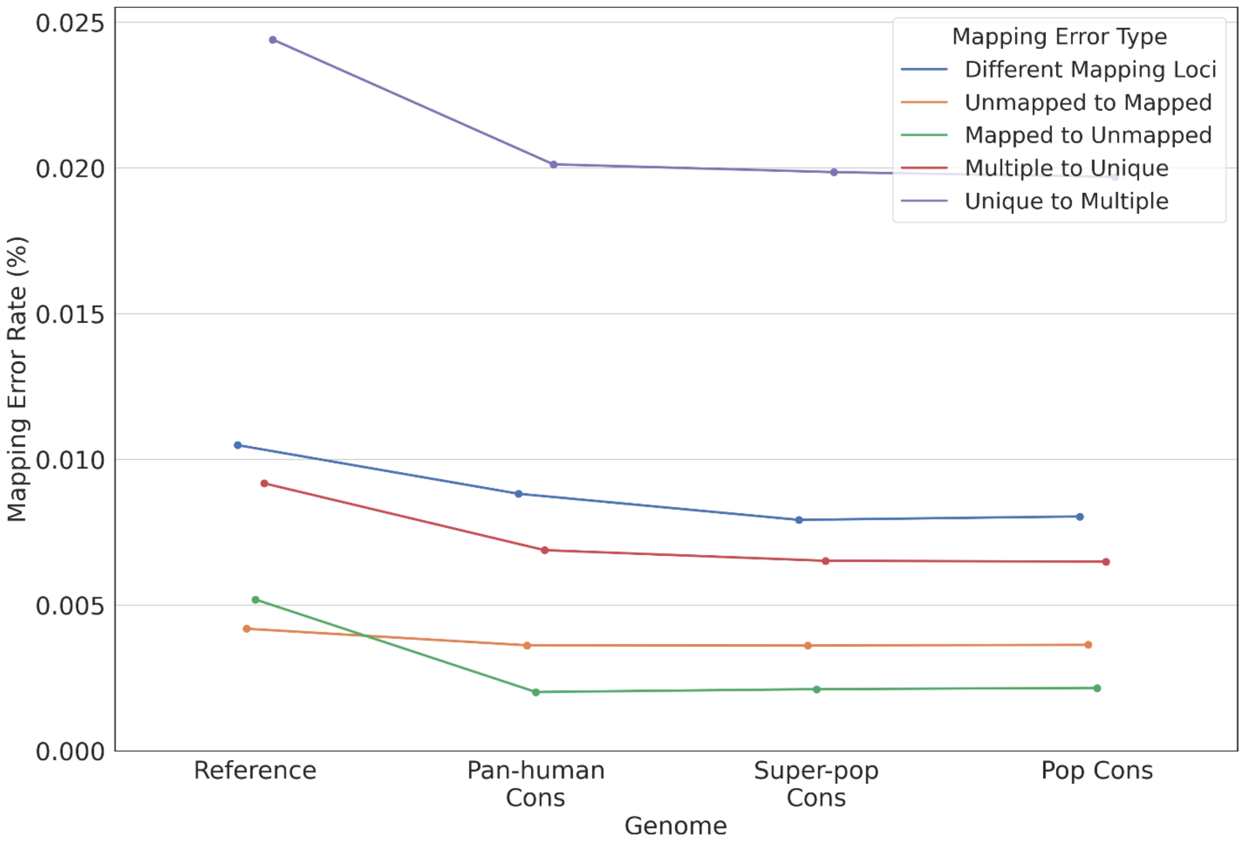

**Supplementary Figure S18:** Overall mapping error rate of reads overlapping insertions or deletions for each error type for individual NA19239.

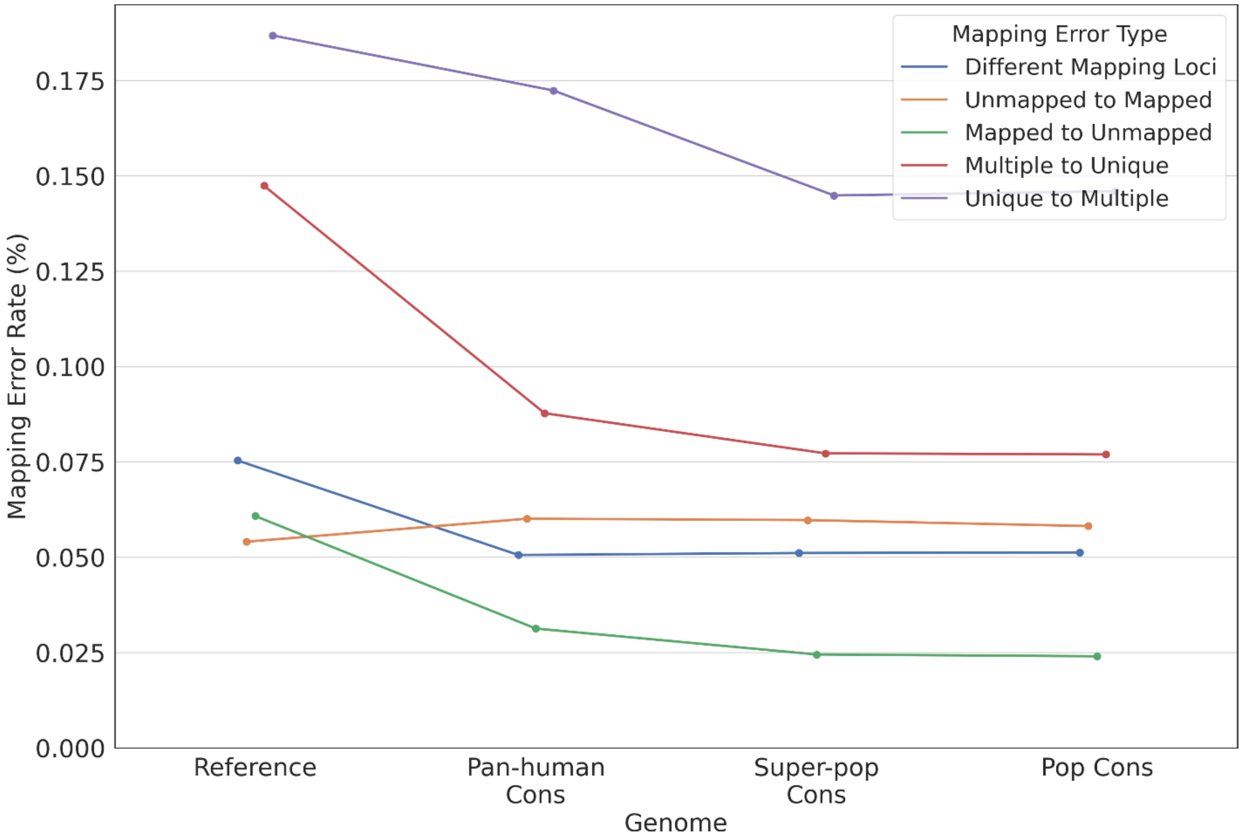

**Supplementary Figure S19:** Overall mapping error rate for each error type for individual NA19240.

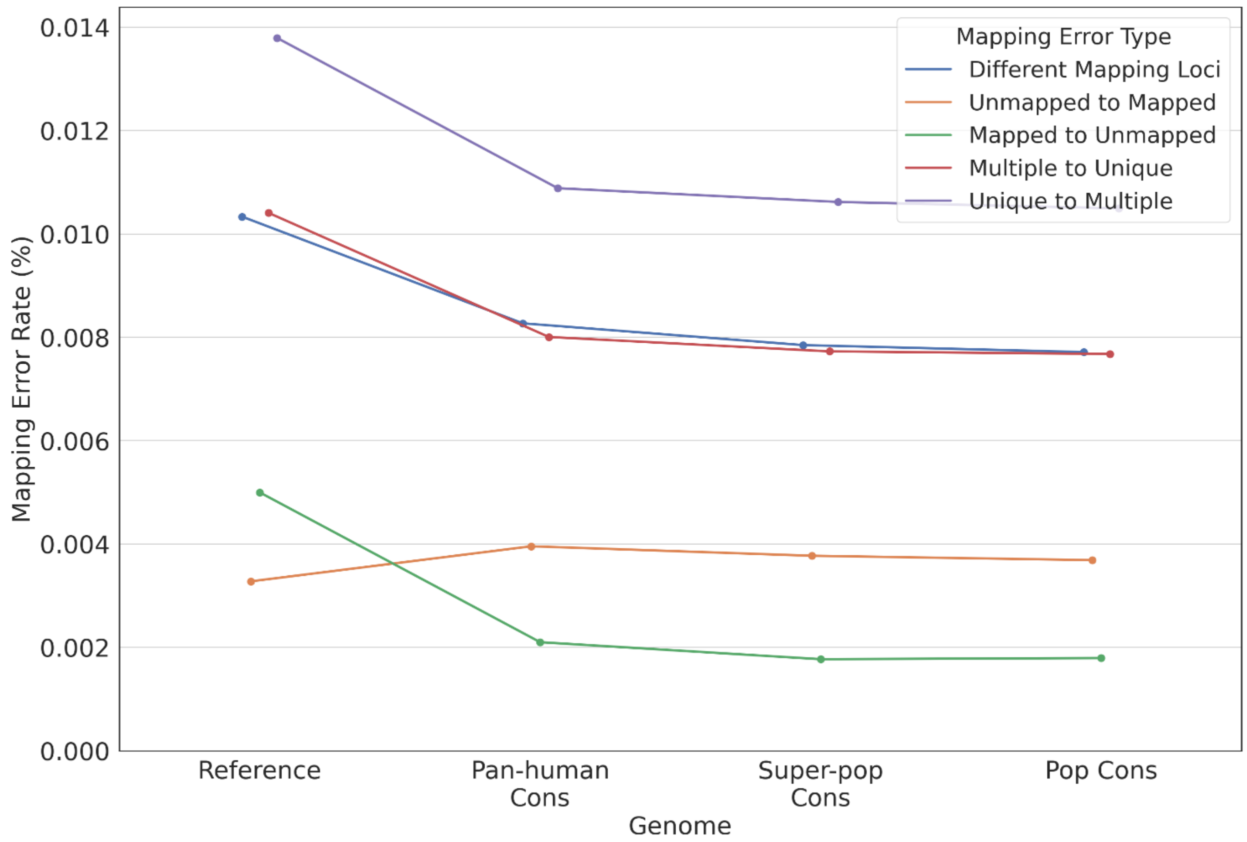

**Supplementary Figure S20:** Overall mapping error rate of reads overlapping insertions or deletions for each error type for individual NA19240.

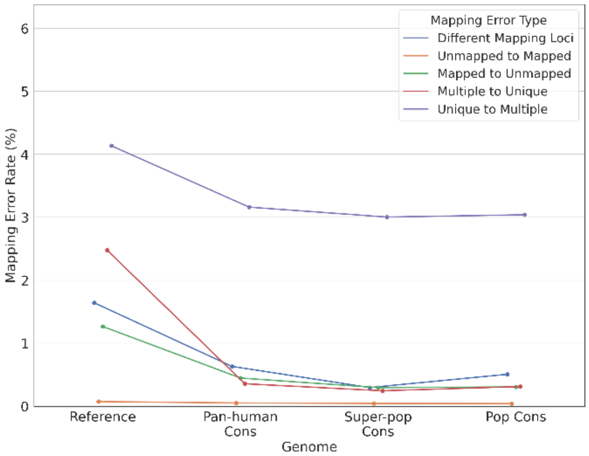

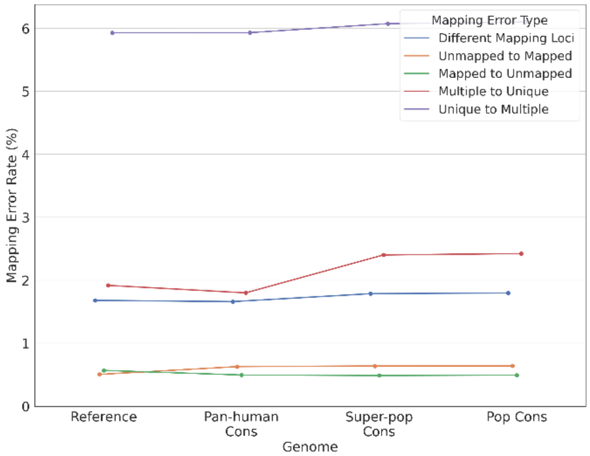

**Supplementary Figure S21:** Homozygous (left) and heterozygous (right) mapping error rate for each error type for individual HG00512.

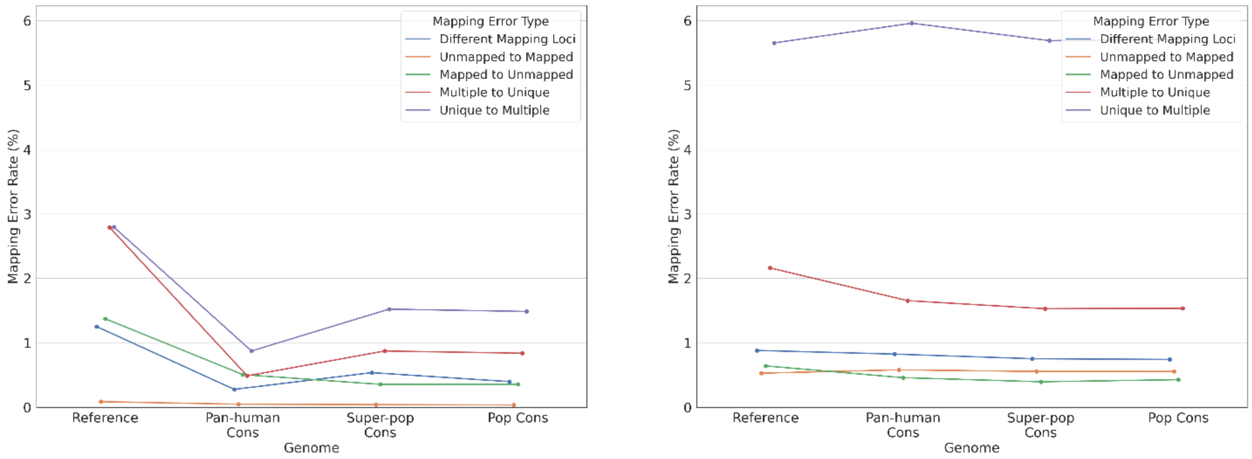

**Supplementary Figure S22:** Homozygous (left) and heterozygous (right) mapping error rate for each error type for individual HG00513.

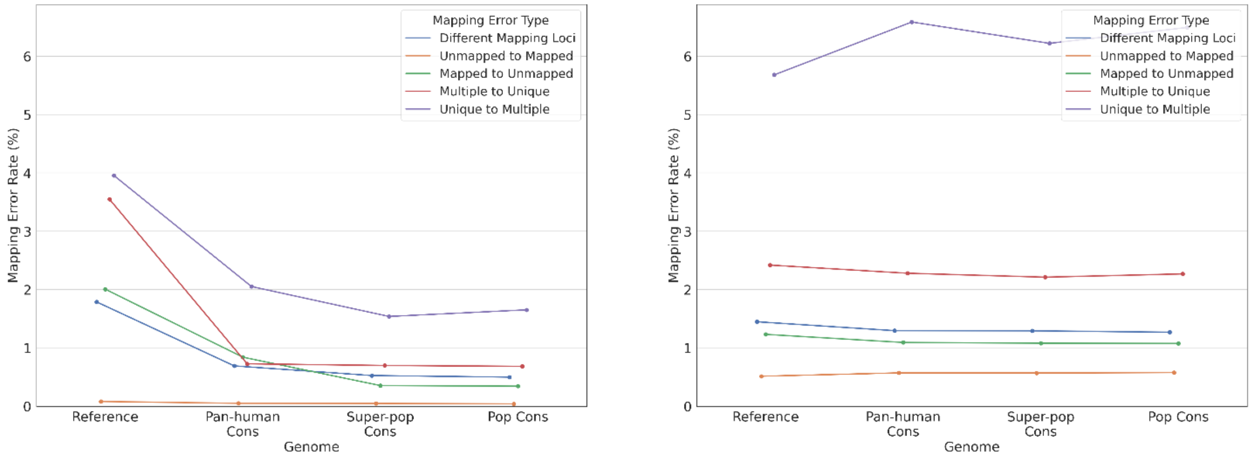

**Supplementary Figure S23:** Homozygous (left) and heterozygous (right) mapping error rate for each error type for individual HG00731.

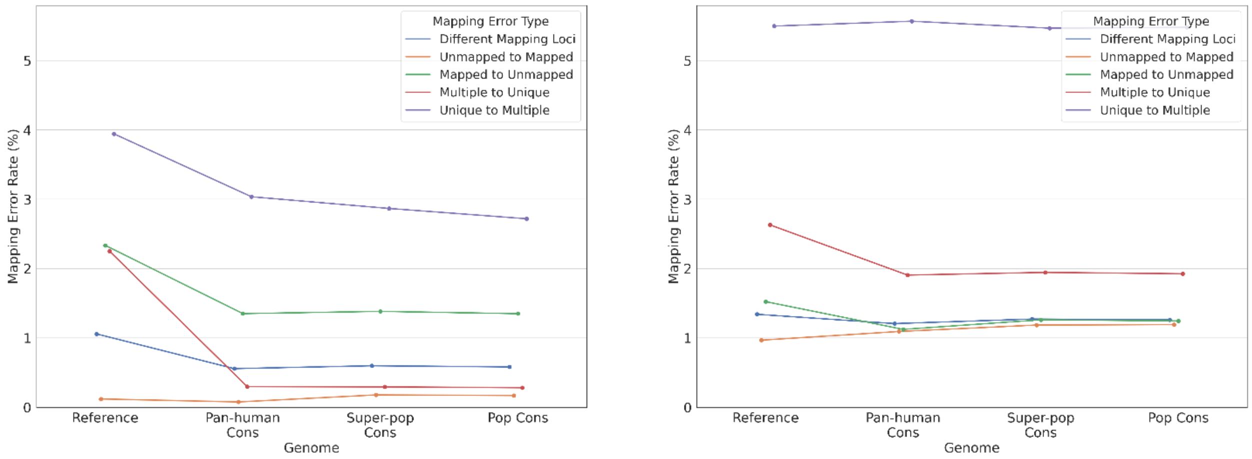

**Supplementary Figure S24:** Homozygous (left) and heterozygous (right) mapping error rate for each error type for individual HG00732.

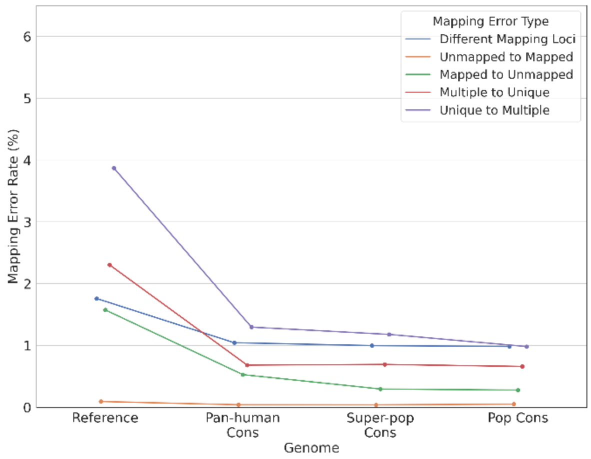

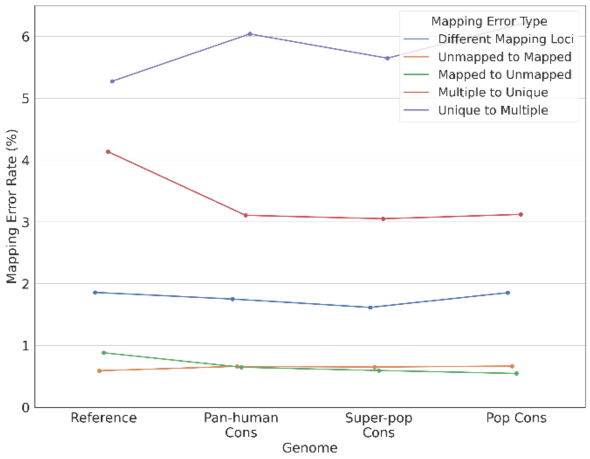

**Supplementary Figure S25:** Homozygous (left) and heterozygous (right) mapping error rate for each error type for individual HG00733.

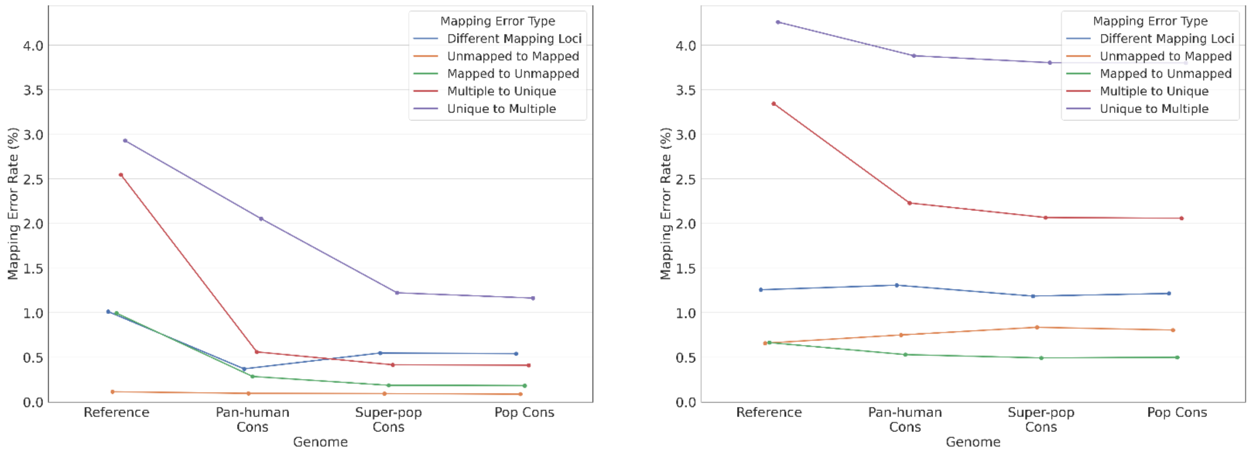

**Supplementary Figure S26:** Homozygous (left) and heterozygous (right) mapping error rate for each error type for individual NA19238.

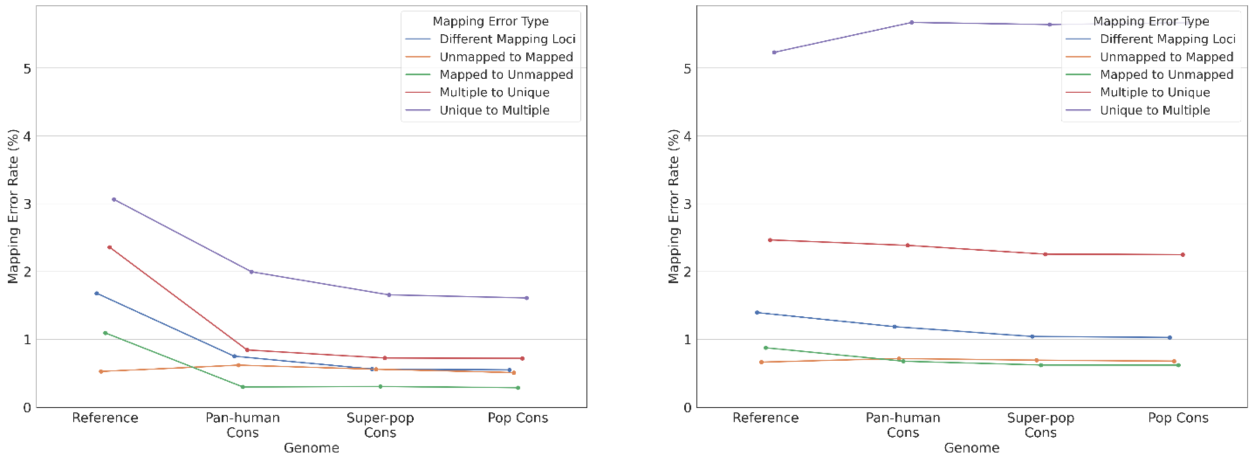

**Supplementary Figure S27:** Homozygous (left) and heterozygous (right) mapping error rate for each error type for individual NA19239.

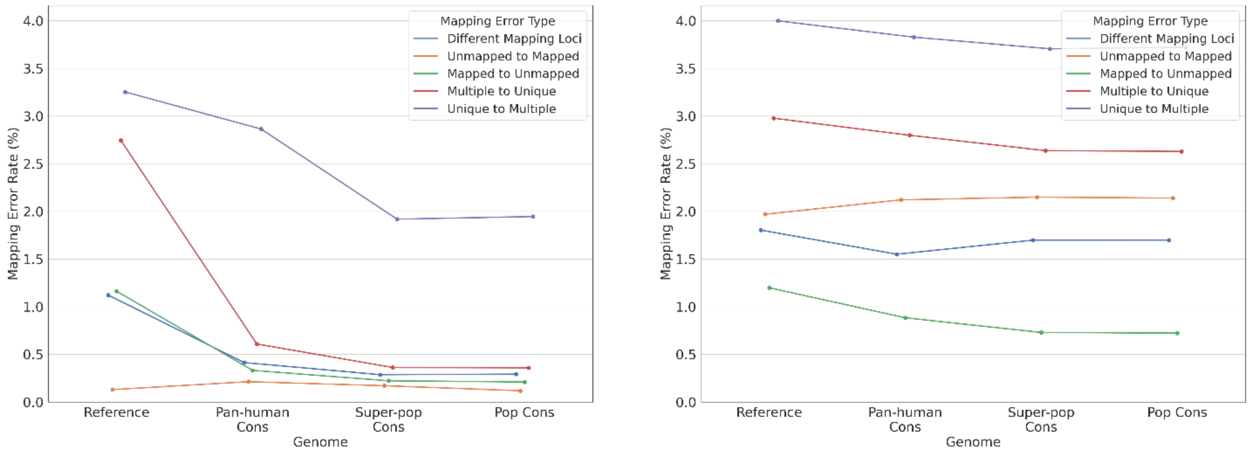

**Supplementary Figure S28:** Homozygous (left) and heterozygous (right) mapping error rate for each error type for individual NA19240.

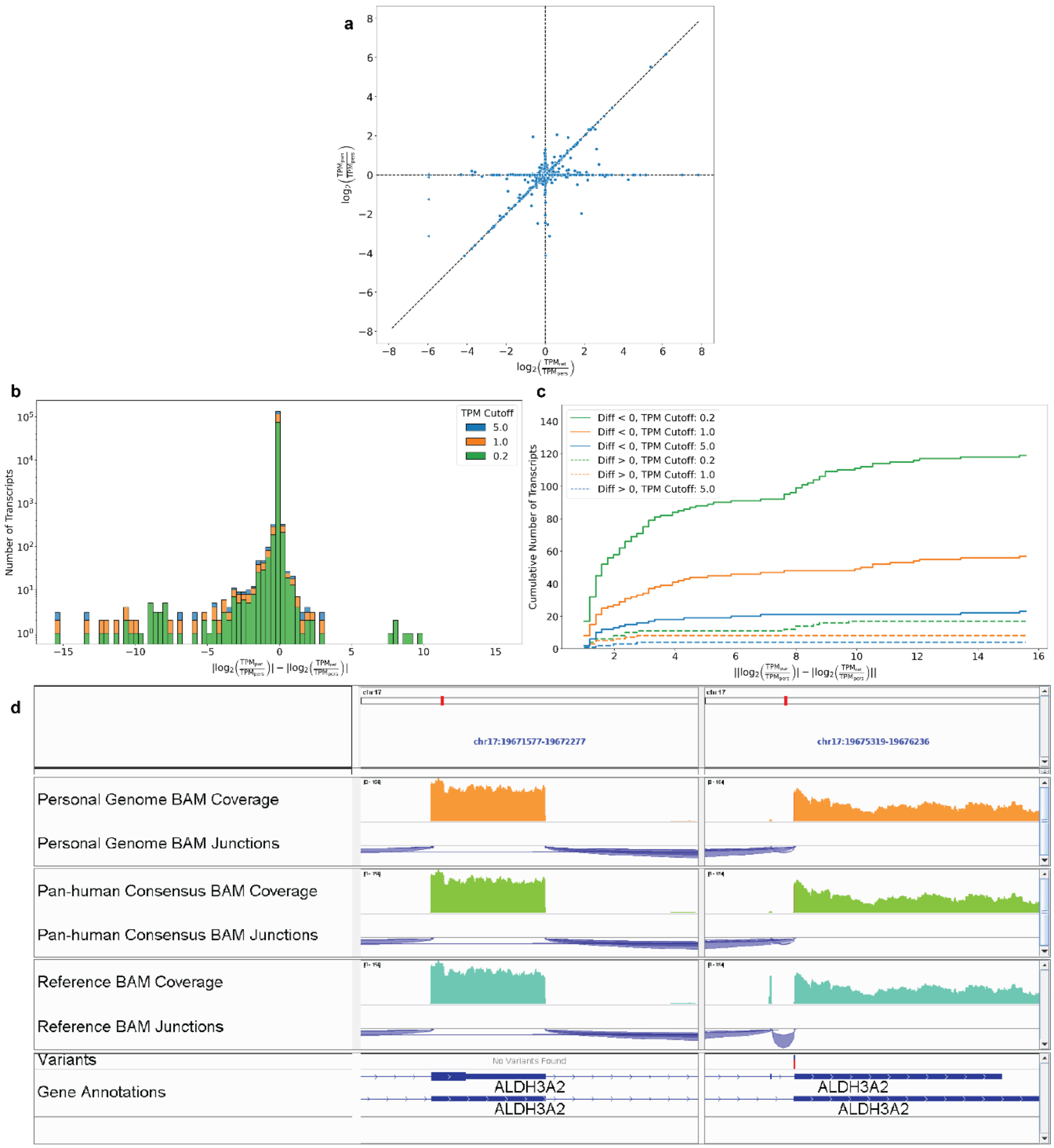

**Supplementary Figure S29: a)** Error in transcript quantification in Pan-human vs Reference genome with respect to the Personal genome. Triangles indicate infinite error (i.e. zero expression in one of the genomes).

**b)** Difference between absolute values of Pan-human to Personal and Reference to Personal log-ratios. Different TPM thresholds are represented by different colors.

**c)** Cumulative distribution of the transcript quantification error. Solid lines represent transcripts which have larger quantification errors in the Reference than in the Pan-human genome; dashed lines represent the opposite cases.

**d)** Read coverage and splice junction tracks for HG00512 reads aligned to the Reference, Pan-human consensus, and HG00512 personal genome. The regions shown are part of the ALDH3A2 gene. The Variants track shows the location of one MAR that is present in the Pan-human consensus and the personal genome.
